# Follicular Bcl6 reactivity is associated with a unique immune landscape and spatial transcriptome in COVID-19

**DOI:** 10.1101/2024.11.07.622471

**Authors:** Cloe Brenna, Bramon Mora Bernat, Kalliopi Ioannidou, Simon Burgermeister, Julien Bodelet, Mia Siebmanns, Spiros Georgakis, Michail Orfanakis, Nazanin Sédille, Matthew J Feinstein, Jon W Lomasney, Oliver Y. Chen, Giuseppe Pantaleo, Sabina Berezowska, Laurence De Leval, Raphael Gottardo, Constantinos Petrovas

## Abstract

The regulation of follicular (F) and germinal center (GC) immune reactivity in human lymph nodes (LNs), particularly during the early stages of viral infections, remains poorly understood. We have analyzed lung-draining lymph nodes (LD-LNs) from COVID-19 autopsies using multiplex imaging and spatial transcriptomics to examine the immune landscape and with respect to the aging. We identified three subgroups of Reactive Follicles (RFs) based on Bcl6 prevalence, RF-Bcl6no/low, RF-Bcl6int and RF-Bcl6high. RF-Bcl6high tissues express a distinct B/TFH immune landscape associated with increased prevalence of proliferating B- and TFH-cell subsets. Comparison between LD-LNs and matched subdiaphragmatic LNs revealed a disconnected Bcl6 reactivity between the two anatomical sites. LD-LNs Bcl6 reactivity was associated with a distinct spatial transcriptomic profile. TH1-associated genes/pathways (e.g. CXCR3, STAT5, TNF signaling) were significantly upregulated in RF-Bcl6no/low tissues while the RF-Bcl6high tissues exhibited significant upregulation of GC-promoting genes/pathways (e.g. CXCL13, B cell receptor signaling). Despite the similar prevalence, the in-situ transcriptome profiling indicates a higher monocyte/macrophage functionality in “Aged” compared to “Young” follicles from donors with comparable Bcl6 reactivity. Our findings reveal a heterogeneous F/GC landscape in COVID-19 LD-LNs and highlight specific molecular targets and pathways that could regulate human F/GC immune dynamics during early viral infections.

## INTRODUCTION

The COVID-19 pandemic, caused by SARS-COV-2, has significantly impacted global health, highlighting the urgent need to understand the human immune responses against the virus(1). Particularly vulnerable groups, such as the elderly and individuals with comorbidities, are at higher risk for severe effects and mortality(2). Age-related immune decline, or “immunosenescence”, further compromises the ability to effectively combat infections and respond to vaccination(3).

Lymph Nodes (LNs) are crucial for shaping immune responses against pathogens(4). With respect to respiratory infections, thoracic LNs, e.g. Hilar Lymph Nodes (HLNs), located in the lung hilum, are of particular interest as they host antigen-presenting cells (APCs) that capture viral antigens from the lungs, facilitating the activation of naïve T-cells into effectors to fight the virus. Within the germinal centers (GCs), the coordinated function of TFH-cells and GC B-cells can lead to the development of pathogen-specific B-cell responses(5). TFH cells assist B cells in undergoing somatic hypermutation (SHM) and affinity maturation, essential for producing high-affinity antibodies and long-term immune memory. Acute COVID-19 infection has been associated with a reduction in GCs and compromised expression of Bcl6^high^ cells, alongside an overexpression of TNF-α that could impair the TFH-cell differentiation and development of GC immune reactivity leading to the generation of long-term B-cell memory(6). This impairment may lead to a compromised response upon reinfection or exposure to new variants(7). The study of relevant lung-draining lymph nodes (LD-LNs) could also provide insights into regional immune responses like effector CD8 T-cells in the lungs, the primary site of SARS-COV-2 infection.

The cellular and molecular mechanisms underlying the development of human GC immune reactivity, particularly in early viral infections, are not well understood. Lack of access to relevant tissues represents a major challenge for such research. Although COVID-19 infection is associated with a highly inflammatory environment in moderate and severe diseases not necessarily found in other viral infections(8), using relevant COVID-19 autopsies could provide useful information regarding basic immunological mechanisms mediating the development of GC reactivity.

The primary aim of our study was to deepen our understanding of GC immune reactivity in the context of COVID-19. To achieve this, we examined LD-LNs from both “Aged” (≥60 years) and “Young” (<57 years) individuals, employing serological measurements, multiplex immunofluorescence imaging (mIF), and spatial *in situ* transcriptomic profiling. This approach allowed us to characterize the cellular and molecular landscape within these LNs during SARS-CoV-2 infection, with a particular focus on how Bcl6 expression relates to the formation of distinct RF profiling. Our findings highlight age-related differences in RFs, with Bcl6 expression corresponding to unique cellular profiles and molecular signatures within the germinal center. Mechanistically, our data suggest that in COVID-19 infection, overexpression of a TH1 follicular signature may hinder the formation of mature RFs, particularly in LD-LNs.

## MATERIALS AND METHODS

### Human material

LN tissues were obtained from the Institute of Pathology at Lausanne University Hospital, Switzerland, and the Feinberg School of Medicine of Northwestern University **(Tables 1 and 2)**. Postmortem examinations and autopsies from patients who died of COVID-19 disease were performed **i)** at the Institute of Pathology of Lausanne University Hospital (CHUV) between March 2020 and March 2021 and **ii)** Department of Pathology, Northwestern University between March 2020, and July 2021. The studies were approved by the Ethical Committee of **i)** the Canton de Vaud, Switzerland (protocol number 2020-01257) and **ii)** Northwestern University (STU# 00202918). Written consent was obtained from all living participants, and in cases where samples were previously donated to tissue repositories from deceased patients, an IRB-approved waiver of consent was applied (Northwestern University, CHUV). Tonsillar tissues were obtained from anonymized children who underwent routine tonsillectomy at the Hospital de l’Enfance of Lausanne, with approval from the Canton de Vaud-CER-VD, Switzerland (PB_2016-02436 (201/11)). This research project was conducted according to the principles of the Declaration of Helsinki.

**Table 1.**
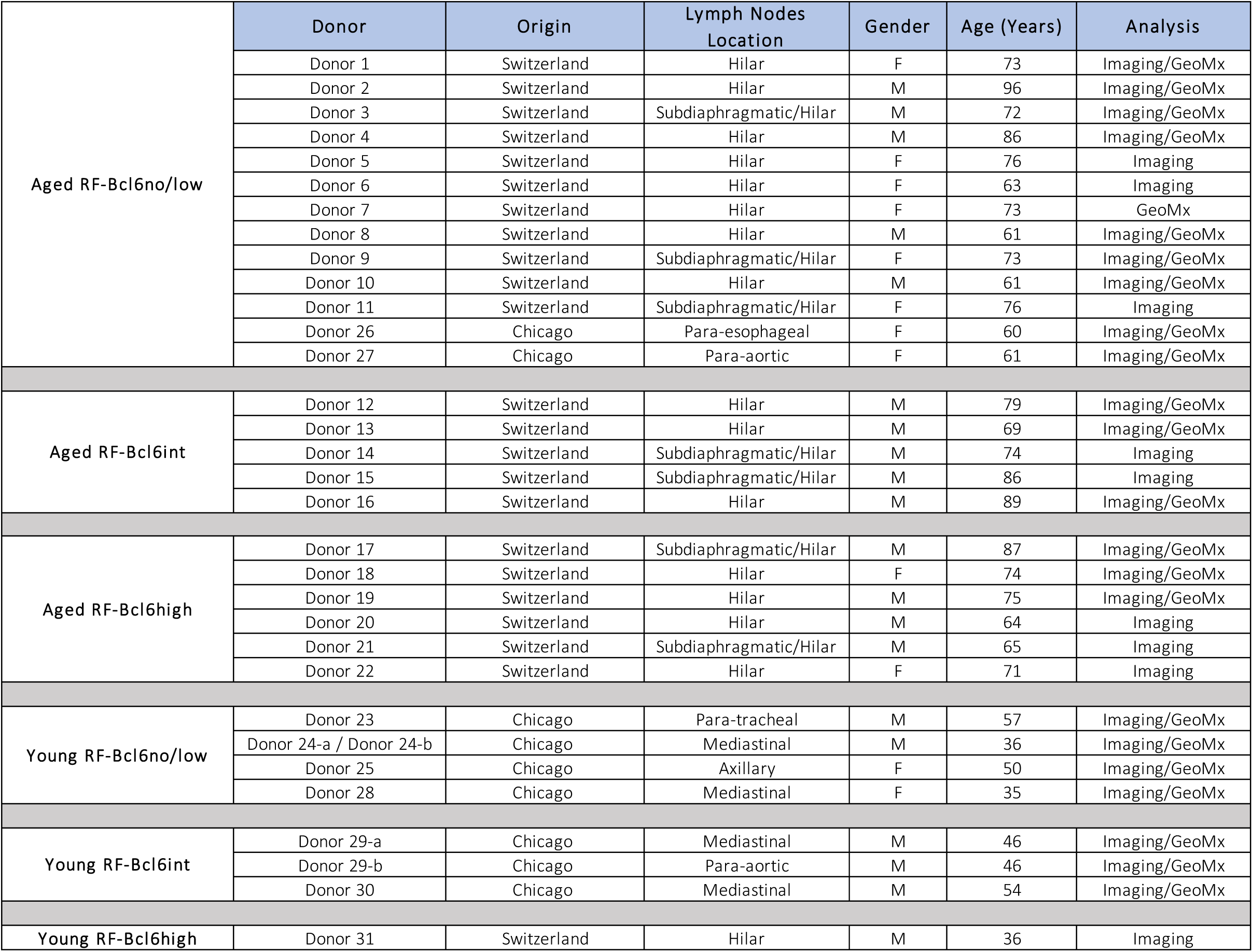
Study Cohort. The origin, anatomical location, demographic data, and the assay applied for each tissue and donor are listed.

|  | Donor | Origin | Lymph Nodes Location | Gender | Age (Years) | Analysis |
| --- | --- | --- | --- | --- | --- | --- |
| Aged RF-Bcl6no/low | Donor 1 | Switzerland | Hilar | F | 73 | Imaging/GeoMx |
|  | Donor 2 | Switzerland | Hilar | M | 96 | Imaging/GeoMx |
|  | Donor 3 | Switzerland | Subdiaphragmatic/Hilar | M | 72 | Imaging/GeoMx |
|  | Donor 4 | Switzerland | Hilar | M | 86 | Imaging/GeoMx |
|  | Donor 5 | Switzerland | Hilar | F | 76 | Imaging |
|  | Donor 6 | Switzerland | Hilar | F | 63 | Imaging |
|  | Donor 7 | Switzerland | Hilar | F | 73 | GeoMx |
|  | Donor 8 | Switzerland | Hilar | M | 61 | Imaging/GeoMx |
|  | Donor 9 | Switzerland | Subdiaphragmatic/Hilar | F | 73 | Imaging/GeoMx |
|  | Donor 10 | Switzerland | Hilar | M | 61 | Imaging/GeoMx |
|  | Donor 11 | Switzerland | Subdiaphragmatic/Hilar | F | 76 | Imaging |
|  | Donor 26 | Chicago | Para-esophageal | F | 60 | Imaging/GeoMx |
|  | Donor 27 | Chicago | Para-aortic | F | 61 | Imaging/GeoMx |
| Aged RF-Bcl6int | Donor 12 | Switzerland | Hilar | M | 79 | Imaging/GeoMx |
|  | Donor 13 | Switzerland | Hilar | M | 69 | Imaging/GeoMx |
|  | Donor 14 | Switzerland | Subdiaphragmatic/Hilar | M | 74 | Imaging |
|  | Donor 15 | Switzerland | Subdiaphragmatic/Hilar | M | 86 | Imaging |
|  | Donor 16 | Switzerland | Hilar | M | 89 | Imaging/GeoMx |
| Aged RF-Bcl6high | Donor 17 | Switzerland | Subdiaphragmatic/Hilar | M | 87 | Imaging/GeoMx |
|  | Donor 18 | Switzerland | Hilar | F | 74 | Imaging/GeoMx |
|  | Donor 19 | Switzerland | Hilar | M | 75 | Imaging/GeoMx |
|  | Donor 20 | Switzerland | Hilar | M | 64 | Imaging |
|  | Donor 21 | Switzerland | Subdiaphragmatic/Hilar | M | 65 | Imaging |
|  | Donor 22 | Switzerland | Hilar | F | 71 | Imaging |
| Young RF-Bcl6no/low | Donor 23 | Chicago | Para-tracheal | M | 57 | Imaging/GeoMx |
|  | Donor 24-a / Donor 24-b | Chicago | Mediastinal | M | 36 | Imaging/GeoMx |
|  | Donor 25 | Chicago | Axillary | F | 50 | Imaging/GeoMx |
|  | Donor 28 | Chicago | Mediastinal | F | 35 | Imaging/GeoMx |
| Young RF-Bcl6int | Donor 29-a | Chicago | Mediastinal | M | 46 | Imaging/GeoMx |
|  | Donor 29-b | Chicago | Para-aortic | M | 46 | Imaging/GeoMx |
|  | Donor 30 | Chicago | Mediastinal | M | 54 | Imaging/GeoMx |
| Young RF-Bcl6high | Donor 31 | Switzerland | Hilar | M | 36 | Imaging |

**Table 2.** Study Cohort. Pulmonary and Immune-Related Comorbidities in the Study.

|  | Donor | Death Post Infection (DPI) | Comorbidities |  |
| --- | --- | --- | --- | --- |
|  |  |  | Pulmonary | Others |
| Aged RF-Bcl6no/low | Donor 1 | 4 | COPD not staged. / Former smoker (58py), stopped 3 years ago. | Bipolar disorder, under lithium treatment. / Substituted hypothyroidism. / Systemic hypertension with peripheral arterial disease. |
|  | Donor 2 | 8 | h/o community acquired pneumonia 1 year ago. / h/o pulmonary embolism (right middle and inferior lobe) 1 year ago. | Severe scoliosis. / Femoral head prosthesis for 2 years. / History of TURP. |
|  | Donor 3 | 1 | Sleep apnea syndrome. | Follicular lymphoma grade I-II (in remission). / Lymphopenia (0.23G/L). / Hypothyroidism (substitution therapy). / Mixed anxiety-depressive disorder (treated with Citalopram). / Cardiomyopathy (arrhythmogenic, hypertensive, and toxic due to chemotherapy). / Systemic hypertension. |
|  | Donor 4 | 19 | - | Cholelithiasis. / Cholecystectomy for cholecystolithiasis. / Minor neurocognitive disorders not investigated. / Myocardial amyloidosis |
|  | Donor 5 | 4 | History of heavy smoking (50 pack-years), currently abstinent. | Large-cell neuroendocrine carcinoma with unknown primary, diagnosed in January 2021, and a history of squamous cell carcinoma of the uvula and soft palate treated in 2012. / Inflammatory anemia related to neoplasia. / Proximal deep vein thrombosis (DVT) in the left lower extremity due to active cancer. / Peripheral artery disease in the lower limbs. / History of alcohol consumption (now abstinent). / Treated hypothyroidism. / Systemic hypertension. |
|  | Donor 6 | - | Pulmonary Langerhans cell histiocytosis (lung transplant in 2004, on immunosuppressants). | Cervico-dorsal myelopathy (inflammatory origin), severe cognitive impairment (related to COVID-19 encephalopathy and possible neurodegenerative disorder). / Treated hypertension and hypercholesterolemia. / Ulcers on both ankles, vertebral fracture at T10 with osteopenia. / Neurogenic bladder with urinary incontinence. / Multiple pancreatic cystic lesions (IPMNs), vulvar condylomas, and cervical dysplasia (treated with conization in 2014). |
|  | Donor 7 | - | - | Alcohol-related liver cirrhosis (CHILD B). / Atrial fibrillation. / Treated hypertension. / Monoclonal B-cell lymphocytosis without palpable lymphadenopathy. / Severe thrombocytopenia. |
|  | Donor 8 | 15 | COPD with asthma component. / Active smoking (80py). | Alcoholism. / Prostate cancer (autopsy finding). / DM type II. / Systemic hypertension with peripheral arterial disease. |
|  | Donor 9 | 14 | - | Squamous cell carcinoma of the lung (autopsy finding). / Paranoid schizophrenia (institutionalized patient). / History of breast carcinoma (8 years ago). / Malnutrition. / Systemic hypertension. |
|  | Donor 10 | - | - | - |
|  | Donor 11 | 55 | Obstructive Sleep Apnea Syndrome (SAOS), managed with CPAP. | Treated hypertension and hypercholesterolemia. / Moderate mitral insufficiency. / Cerebral aneurysm rupture in 1990 with placement of a ventricular drain. / Right brachio-crural motor hemiparesis of indeterminate origin. / Vascular encephalopathy. |
|  | Donor 26 | 39 | Chronic Obstructive Pulmonary Disease (COPD). / Bilateral lung transplant (3 years prior). | Heart failure. / Diabetes. |
|  | Donor 27 | 17 | - | Hypertension. / Diabetes. |
| Aged RF-Bcl6int | Donor 12 | 12 | COPD stage II. / Asthma. | Prostate cancer (autopsy finding). / Pancreatic neuroendocrine tumor (autopsy finding). / Ankylosing spondylarthritis. / History of cerebellar ischemic stroke (9 years ago). / DM type II. / Dyslipidemia. / AVB 1st degree and LAFB. |
|  | Donor 13 | - | - | Dilated cardiomyopathy (mixed: ischemic, hypertensive, genetic). / History of coronary stenting. / Automatic defibrillator (for 9 years). / Systemic hypertension. / Dyslipidemia. |
|  | Donor 14 | 38 | - | High-grade papillary urothelial carcinoma (pT2b pN0 R0 M0) with Bricker urinary diversion. / Hypertensive heart disease. / Treated hypertension. / Type 2 diabetes (non-insulin-dependent, treated). / Overweight (BMI 29 kg/m<sup>2</sup>). / Treated dyslipidemia. |
|  | Donor 15 | 1 | - | Second-degree atrioventricular block (AV block). / Myocardial amyloidosis. / Severe coronary artery disease. / Treated hypertension. / Alzheimer's-type degenerative encephalopathy, Braak stage 3. / Cerebro-meningeal amyloid angiopathy. / Osteoporosis. |
|  | **Donor 16** | - | - | Heart failure with preserved EF, managed with pacemaker and conservative treatment for coronary issues. / Treated hypertension. / History of prostate and joint surgeries (hip and shoulder). / Multifactorial balance issues, mild neurocognitive decline, and suspected lacunar stroke. / Anemia, mild Amiodarone-induced hyperthyroidism, and previous drug-induced skin reaction. |
| **Aged RF-Bcl6high** | **Donor 17** | 4 | - | DM type II. / Dyslipidemia. |
|  | **Donor 18** | 9 | COPD stage III. / Asthma. / Sleep apnea syndrome. | Mixed anxiety-depressive disorder (treated). / Hiatal hernia. / Substituted hypothyroidism following subtotal thyroidectomy for multinodular goiter. / Arrhythmogenic cardiomyopathy (pacemaker). / Systemic hypertension. |
|  | **Donor 19** | 23 | Former smoker. | **Metatarso-phalangeal osteoarthritis (left hallux) due to diabetic foot ulcer with** secondary bacteremia (methicillin-susceptible Staphylococcus aureus and Proteus mirabilis) (1 month before death). / DM type II. / Hypertrophic and valvular cardiomyopathy. / History of aortic valve replacement. / Pacemaker due to complete heart block. / Systemic hypertension. |
|  | **Donor 20** | 11 | - | Treated hypertension (Exforge). / Liver steatosis, colonic polyps (2014, 2018), and gastroesophageal reflux (2001). / Suspected right ureteropelvic junction disorder and history of ureteral stone (1989). / Dupuytren's disease. |
|  | **Donor 21** | 16 | Neuroendocrine small-cell lung carcinoma with bone and liver metastases, diagnosed in September 2019. Under palliative treatment. / Former smoker (50 pack-years), quit in 2016. | Left foot drop likely due to peripheral involvement. / Coronary artery disease with NSTEMI in December 2016 and stent placement in the marginal and right coronary arteries. |
|  | **Donor 22** | 13 | - | Regenerative normocytic normochromic anemia. / Acute confusional state (hypo and hyperactive) of multifactorial origin. / Gait and balance instability. / Mixed anxiety-depressive disorder. / Moderate mental retardation. / History of hysterectomy (not dated). / Ischemic cardiomyopathy. / History of myocardial infarction (not dated). / History of myocarditis 8 years ago. |
| **Young RF-Bcl6no/low** | **Donor 23** | 20 |  | GE reflux. / Gout. / Seizure disorder. / Gunshot wound 1985 (head and chest). / Hypertension. / Stroke. / Hyperlipidemia. |
|  | **Donor 24-a / Donor 24-b** | 40 | - | Obesity. / Diabetes. / Central Hypothyroidism. / Hypertension. / Hyperlipidemia. |
|  | **Donor 25** | 30 | - | Alcoholic Cirrhosis. / Hepatic Encephalopathy. / Obesity. / Gastric Bypass surgery. |
|  | **Donor 28** | 50 | Asthma. | Obesity. / Systemic Lupus Erythematosus. / Sickle cell trait. |
| **Young RF-Bcl6int** | **Donor 29-a / Donor 29-b** | 47 | - | Obesity. |
|  | **Donor 30** | 27 | - | Hyperlipidemia. |
| **Young RF-Bcl6high** | **Donor 31** | 14 | - | Tetraplegia (due to a road accident 13 years ago). |

#### Sex as a biological variable

Our study examining both male and female participants. Due to the small sample size, sex was not considered as a covariate in this investigation. Our cohort comprised 20 males (14 in the "Aged" group and 6 in the "Young" group) and 12 females (10 in the "Aged" group and 2 in the "Young" group).

Plasma was collected from some of the participants using EDTA as an anticoagulant. An immunoturbidimetric assay was employed for the quantification of CRP levels. Absolute numbers of leukocytes (G/L), lymphocytes (G/L), neutrophils (G/L), monocytes (G/L), eosinophils (G/L), and basophils (G/L) were determined using an automated hematology analyzer.

### Multiplex Immunofluorescent (mIF)-data acquisition

4µm thick sections were prepared from formalin-fixed, paraffin-embedded (FFPE) blocks and mounted on Superfrost glass slides (Thermo Scientific, Waltham). The sections were then used for mIF and spatial transcriptomic (GeoMx) analysis. The tissue staining involved several sequential steps as previously described(9). Antigen retrieval was followed by blocking of non-specific antibody binding using the Opal blocking/antibody diluent solution. Then, titrated primary antibodies and appropriate secondary HRP-labeled antibodies were applied **(Tables 3 and 4)**. The detection of the target proteins was achieved using optimized fluorescent Opal tyramide signal amplification (TSA) dyes (Akoya), along with repeated cycles of antibody denaturation. To visualize the cell nuclei, the sections were counterstained with Spectral DAPI. Following staining, the sections were rinsed in water with soap and mounted using DAKO mounting medium from Dako/Agilent, Santa Clara. Multispectral images (MSI) were acquired using **i)** the Vectra Polaris 1.0 imaging system (Akoya) at a resolution of 5µm/pixel (20x) and **ii)** a Leica Stellaris 8 SP8 confocal system, equipped with LAS-X software (at 512 × 512-pixel density and 20× objective). A compensation matrix, which was created using the Leica LAS-AF Channel Dye Separation module (Leica Microsystems) and tissue sections stained with a single antibody–fluorophore, was used to correct fluorophore spillover across different channels.

**Table 3.** List of primary antibodies used in the study. The origin source (host), clone, isotype and vendor are provided.

| *Primary Antibodies* | *Host* | *Clonality* | *Clone* | *Isotype* | *Vendor* | *Cat #* |
| --- | --- | --- | --- | --- | --- | --- |
| PD1 | mouse | monoclonal | NAT 105 | IgG1k | Biocare medical | ACI 3137 |
| CD57 | mouse | monoclonal | NK-1 | IGMk | Diagnostic BioSystem | Mob 163 |
| Bcl-6 | mouse | monoclonal | GI191E/A8 | IgG | CELL MARQUE | 760-4241 |
| CD20 | mouse | monoclonal | L26 | IgG1k | Novocastra-Leica | NCL-L-CD20-L26 |
| CD4 | rabbit | monoclonal | SP35 | IgG | VENTANA MEDICAL | 790-4423 |
| Ki67 | mouse | monoclonal | MIB-1 | IgG1k | DAKO | M7240 |
| GrzB | mouse | monoclonal | GrB-7 | IgG2a | MONOSAN | MON7029C |
| CD8 | mouse | monoclonal | C8/144B | IgG1k | DAKO | M7103 |
| CD14 | rabbit | monoclonal | EPR3653 | IgG | CELL MARQUE | 114R-15 |
| MPO | rabbit | polyclonal | EPR4793 | IgG | abcam | AB9535 |
| CD68 | mouse | monoclonal | PG-M1 | IgG3k | DAKO | M0876 |
| FDC | mouse | monoclonal | CNA.42 | IGM | Invitrogen | 14-9968-82 |

**Table 4.** The multiplex panels applied are shown. The primary antibodies /opal fluorochromes and the application order of all primary antibodies used in a given panel are shown.

| Panel | Order | Primary Antibody | Dilution | Opal Fluorophore | Dilution |
| --- | --- | --- | --- | --- | --- |
| Adaptive Immunity | 1 | PD1 | 1/100 | opal 620 | 1/150 |
|  | 2 | CD57 | 1/300 | opal 570 | 1/150 |
|  | 3 | Bcl-6 | dispenser | opal 480 | 1/150 |
|  | 4 | CD20 | 1/400 | opal 520 | 1/150 |
|  | 5 | CD4 | 1/200 | opal 690 | 1/200 |
|  | 6 | Ki67 | 1/300 | opal 780 | 1/25 |
| Innate Immunity | 1 | GzrB | 1/40 | opal 570 | 1/150 |
|  | 2 | CD8 | 1/100 | opal 690 | 1/150 |
|  | 3 | CD20 | 1/400 | opal 480 | 1/700 |
|  | 4 | CD14 | 1/50 | opal 620 | 1/150 |
|  | 5 | MPO | 1/150 | opal 520 | 1/150 |
|  | 6 | CD68 | 1/200 | opal 780 | 1/25 |
| FDC panel | 1 | FDC | 1/200 | alexa 488 | 1/150 |
|  | 4 | CD20 | 1/45 | conj-ef650 | 1/150 |

**Table 5.**
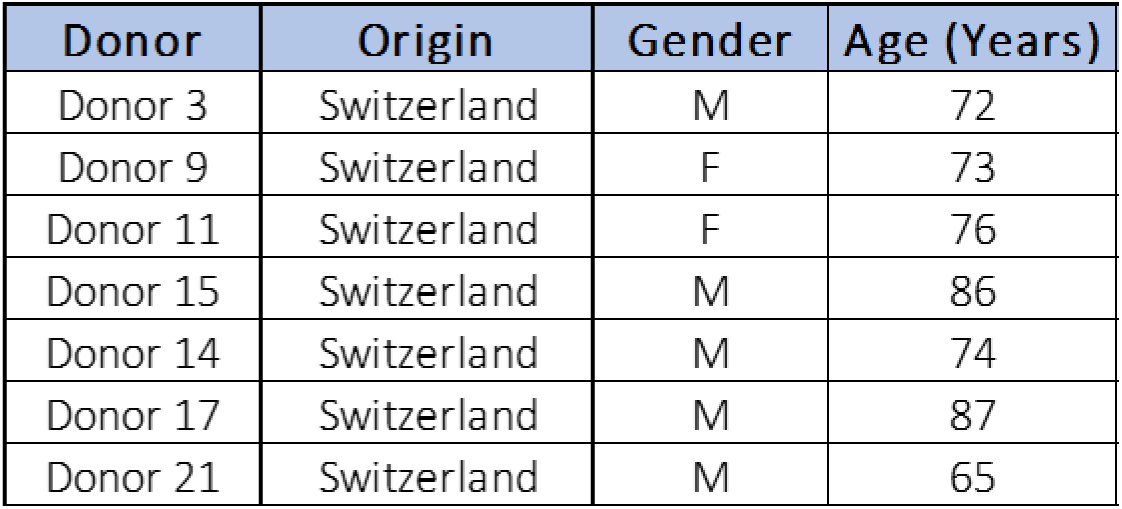
Donors used for the analysis of GC reactivity in Hilar vs. matched control Subdiaphragmatic LNs.

| Donor | Origin | Gender | Age (Years) |
| --- | --- | --- | --- |
| Donor 3 | Switzerland | M | 72 |
| Donor 9 | Switzerland | F | 73 |
| Donor 11 | Switzerland | F | 76 |
| Donor 15 | Switzerland | M | 86 |
| Donor 14 | Switzerland | M | 74 |
| Donor 17 | Switzerland | M | 87 |
| Donor 21 | Switzerland | M | 65 |

### Imaging data analysis

For the Vectra Polaris-generated images, the Phenochart 1.0.12 software (Akoya), a whole-slide viewer for high-resolution multispectral acquisition and annotation capabilities, was used to navigate among slides and identify specific Regions of Interest (ROIs). The subsequent imaging and analysis were performed on the whole tissue. The acquired MSI were analyzed using the InForm analysis software, version 2.4.8 (Akoya). Initially, the images were unmixed, and specific ROIs were used for the algorithm training and cell segmentation across the imaged tissue. The segmentation process involved utilizing the appropriate training components, such as CD20, BCL6, and PD1 to define the GC, Extrafollicular area (EF), and region without tissue. Tissue segmentation was based on selected training markers and DAPI expression, extracting the autofluorescence signal of the tissue. This was achieved by manually drawing training regions for each analyzed image. The segmentation algorithm was trained individually on each image, ensuring a training accuracy of over 90% for the ROIs segmentation. Next, individual cells were segmented using an adaptive cell segmentation algorithm based on the counterstained sections. Cytoplasm and membrane markers were utilized to help in this cell segmentation process. A file report was generated, containing the spatial coordinates (X, Y) of each segmented cell, along with their mean intensity for each Opal fluorophore used. This file report was then extracted and converted into a FACS file (FSC) format before being uploaded to the FlowJo 10 software and the generated data were further analyzed with HistoFlowCytometry(10). Data are reported as normalized numbers per µm2 or frequency of total imaged cells.

### Morphological characteristics of follicular areas

Morphological analysis of individual follicular ROIs, identified based on the density of CD20^high/dim^ B cells, was carried out using FIJI software, where various features such as the area, circularity, and solidity were extracted. To ensure robust analysis, approximately 20 ROIs per tissue were meticulously chosen, each representing distinct and well-defined follicular structures within the tissues. Subsequently, from each ROI, the follicle areas were extracted and normalized using the tissue scale bar. This normalization process ensured that the follicle areas were accurately represented relative to the tissue dimensions, facilitating meaningful comparisons across samples.

### Distance Analysis and Spatial Distribution

To investigate the spatial relationship between relevant cell types (GC-B and TFH-cell subsets), the minimum distance between individual B-cells and TFH-cells, within follicles expressing more than 20 positive cells for each population, was calculated with Python 3.10.9 using the Scipy library(11). The matrix interaction was generated using X and Y coordinates from each cell type and the median distance was extracted. The probability of observing different patterns of cellular distribution across individual ROIs and patients was investigated by analyzing the curves generated from the Ripley’s G function and the theoretical Poisson curve using Pointpats 2.3.0(12). The area between the empirical and theoretical Poisson curve was extracted using the NumPy library(13). The data were presented either as bar graphs (range of all various distances measured x-axis vs frequency or count of B-cells within each distance range y-axis) or dot plots, with each dot representing the mean value of the minimum Euclidean distances between two cell populations for each follicular area (ROI).

### Tissue Spatial Transcriptomic analysis

Transcriptomic profiling was performed using the commercially available platform GeoMx Digital Spatial Profiling (Nanostring). 4 µm FFPE tissue sections from n= 8 RF-Bcl6no/low “Aged” and n=7 RF-Bcl6no/low “Young”, n=3 RF-Bcl6int “Aged” and n=3 RF-Bcl6int “Young”, and n=3 RF-Bcl6high “Aged”, were used. Follicular Regions of Interest (ROIs, n=9-12 per slide) were identified based on the corresponding mIF image and the CD3, CD20 *in situ* staining pattern before the probe-hybridization step. ROIs were selected within a range of areas, and quantitative analysis revealed an average area of 245,745.52µm^2^ (± 211,116.16 µm², SD). The data were collected in two different experiments (A and B). The DSP GeoMx software was used for the PCA analysis of specific gene sets among the section ROIs.

#### Data Processing and Quality Control

We analyzed NanoString DSP data using standard GeoMx processing workflows(14), and R version 4.3.2. Specifically, we processed the DCC files and conducted quality control at the segment, probe, and gene levels using the R packages “GeoMxWorkflows”(14), “GeomxTools”(15), and “NanoStringNCTools”(16). First, we adjusted all zero expression counts to one to enable subsequent data transformations. We then implemented several quality control metrics recommended by NanoString for our segments, including a minimum of 1000 reads, 80% trimming, stitching, and alignment, 50% sequencing saturation, a minimum negative control count of 1, a maximum of 1000 reads observed in NTC wells, and a minimum area of 1000. Next, we removed probes for which the average count across segments was less than 10% of the average count for all probes targeting the same gene across segments, as well as probes that were outliers in at least 20% of the segments. Finally, we filtered out segments and genes with low signal, specifically removing segments where less than 5% of panel genes were detected above the level of quantification (LOQ, defined as two standard deviations above the mean) and genes detected below the LOQ in at least 10% of segments.

#### Batch Correction

to perform batch correction, we first normalized the raw data using the Trimmed Mean of M-values (TMM) method with the R package “standR”(17), which adjusts for differences in library sizes and composition between RNA-seq samples. We then applied the RUV-4 correction from the same package(18), which removes unwanted variations by identifying negative control genes and calculating scaling factors for batch correction. In our analysis, we identified 300 negative control genes and set the number of scaling factors to 2.

#### Differential Gene Expression and Enrichment Analysis

to perform differential gene expression, we followed the limma-voom pipeline(19). We defined a linear model with a design matrix containing the treatment variable and weight matrix from the RUV-4 correction method as covariates. Using this model, we then compared every pair of treatments, identifying differentially expressed genes as those presenting an adjusted p-value smaller than 0.05. Alternatively, we also studied the differences across the different treatments using the R package DESeq2(20), comparing the intercepts for every treatment in a negative binomial regression also accounting for the scaling factors from RUV-4 model. Using the differentially expressed genes in each comparison, we then performed enrichment analysis (GSEA) to identify relevant biological pathways(21). To do so, we employed the “fry” method from the R package limma(19), using the canonical pathways gene sets from the Reactome pathway database(22). The results were then analyzed and visualized using the R package vissE(23).

### Projection Approach for Data Visualization

We used the Statistical Quantile Learning (SQL) method(24), a tool for nonlinear dimensionality reduction, to summarize, analyze, and visualize adaptive and innate immunity data separately. We chose SQL due to its capability to handle high-dimensional datasets and accurately capture complex, nonlinear relationships often found in biological data. The adaptive immune panel featured cell subsets such as CD20^high^, CD20^high^Ki67^high^, CD20^high^Ki67^high^Bcl6^high^, CD4^high^, PD1^high^, PD1^high^Ki67^high^, PD1^high^CD57^high^, and PD1^high^Ki67^high^Bcl6^high^. On the other hand, the innate immune panel included cell subsets like CD8^high^, CD8^high^GrzB^high^, CD14^high^, MPO^high^, and CD68^high^. These markers were specifically chosen to aid in clustering and classifying our immune data and extracted from our Histocytometry analysis. The SQL method provides a straightforward approach to estimating nonlinear latent variable models, or generative models. Unlike local methods such as UMAP, SQL assumes a probabilistic model and learns a generator, which is a smooth function that connects the laten space to the data space. This approach enables SQL to learn a global latent space that captures the overall structure of the data, whereas UMAP emphasizes preventing local neighborhood relationships. The generator allows for easy reconstruction of the data from the latent space, which permits interpretability of the latent space. Compared to other generative methods (such as Variational Autoencoders) SQL is not only easy to fit but also performs better for small samples and large-dimensional data. Additionally, SQL is supported by statistical guarantees and delivers unique and interpretable latent variables. This approach may therefore provide a reliable means to uncover insights that might be overlooked by other methods. Our goal was to cluster our three groups —RF-Bcl6no/low, RF-Bcl6int, and RF-Bcl6high— while also considering different age groups (“Aged” and “Young”). This methodology allowed for clearer separations and deeper insights into adaptive and innate immunity data across varied age groups and offers more meaningful insights, especially in understanding immune response variations within the specified groups.

For the spatial transcriptomic data, Principal Component Analysis (PCA) was employed to analyze specific gene sets across the section ROIs. This technique reduces the high-dimensional gene expression data into principal components that capture the most significant variance, facilitating a clearer visualization of patterns across the different ROI groups.

### Statistical Analysis

Unpaired Non-Parametric Mann-Whitney t-test and Wilcoxon t-test were used for the analysis of mIF data, while a Mixed Effect Model was applied to the serological measurements, comprising fixed effects corresponding to various serological measurements and random effects originating from individual donors. Mean and standard deviation were calculated for normally distributed data, while median and interquartile range were reported for non-normally distributed data. A p value of less than 0.05 was considered significant, with significance levels indicated as follows: p> 0.05 (ns), p ≤ 0.05 (*), p *≤ 0.01 (*\*\*), p ≤ 0.001 (***), and p ≤ 0.0001 (****). Error bars on the graphs represent the mean with the standard error of the mean (SEM). FDR was applied for p value correction, with raw p values and FDR-adjusted p values, along with their significance symbols, reported for the main figures and for the supplementary figures **(Tables 6 and 7)**. The p values displayed on the graphs are derived from the raw P values. Graphs were generated using GraphPad Prism (version 8.3.0), Python (version 3.10.9), and R (version 4.3.0).

**Table 6.** Summary of Statistical Analysis for Cell Population Comparisons: Raw and FDR-adjusted P-values with Significance Indicators Across Multiple Main Figures.

| Figure | Cell Population | Comparison Group | Comparison | Raw p-value | Significance symbol<br>Raw p-value | FDR-adjusted p-value | Significance symbol<br>FDR-adjusted p-value |
| --- | --- | --- | --- | --- | --- | --- | --- |
| Figure 1C | CD20hi Ki67hibcl6hi | Aged | GC-Bcl6no/low vs GC-Bcl6high | 0.0001 | *** | 0.0006 | *** |
|  |  |  | GC-Bcl6no/low vs GC-Bcl6int | 0.0006 | *** | 0.0018 | *** |
|  |  |  | GC-Bcl6int vs GC-Bcl6high | 0.0043 | ** | 0.0086 | ** |
|  |  | Young | GC-Bcl6no/low vs GC-Bcl6int | 0.0536 | ns | 0.0804 | ns |
|  |  | Aged vs Young | Aged GC-Bcl6no/low vs Young GC-Bcl6no/low | 0.9391 | ns | 0.9391 | ns |
|  |  |  | Aged GC-Bcl6int vs Young GC-Bcl6int | 0.0714 | ns | 0.08568 | ns |
| Figure 1D | Normalized Follicle Areas (µm2) | x | GC-Bcl6no/low vs GC-Bcl6high | <0.0001 | **** | 0.00015 | **** |
|  |  |  | GC-Bcl6no/low vs GC-Bcl6int | <0.0001 | **** | 0.00015 | **** |
|  |  |  | GC-Bcl6int vs GC-Bcl6high | 0.115 | ns | 0.115 | ns |
| Figure 1E | CD20hi - count per (µm2) | Aged | GC-Bcl6no/low vs GC-Bcl6high | 0.8201 | ns | 0.8201 | ns |
|  |  |  | GC-Bcl6no/low vs GC-Bcl6int | 0.6461 | ns | 0.8201 | ns |
|  |  |  | GC-Bcl6int vs GC-Bcl6high | 0.7922 | ns | 0.8201 | ns |
|  |  | Young | GC-Bcl6no/low vs GC-Bcl6int | 0.0357 | * | 0.2142 | ns |
|  |  | Aged vs Young | Aged GC-Bcl6no/low vs Young GC-Bcl6no/low | 0.2343 | ns | 0.4686 | ns |
|  |  |  | Aged GC-Bcl6int vs Young GC-Bcl6int | 0.0714 | ns | 0.2142 | ns |
|  | CD20hiKi67hi - count per (µm2) | Aged | GC-Bcl6no/low vs GC-Bcl6high | 0.0047 | ** | 0.0282 | ** |
|  |  |  | GC-Bcl6no/low vs GC-Bcl6int | 0.0194 | * | 0.0582 | * |
|  |  |  | GC-Bcl6int vs GC-Bcl6high | 0.4286 | ns | 0.51432 | ns |
|  |  | Young | GC-Bcl6no/low vs GC-Bcl6int | 0.1429 | ns | 0.2858 | ns |
|  |  | Aged vs Young | Aged GC-Bcl6no/low vs Young GC-Bcl6no/low | 0.9593 | ns | 0.9593 | ns |
|  |  |  | Aged GC-Bcl6int vs Young GC-Bcl6int | 0.25 | ns | 0.375 | ns |
|  | CD20hiKi67hiBcl6hi - count per (µm2) | Aged | GC-Bcl6no/low vs GC-Bcl6high | 0.0001 | *** | 0.0006 | *** |
|  |  |  | GC-Bcl6no/low vs GC-Bcl6int | 0.0023 | ** | 0.0006 | ** |
|  |  |  | GC-Bcl6int vs GC-Bcl6high | 0.0519 | ns | 0.07785 | ns |
|  |  | Young | GC-Bcl6no/low vs GC-Bcl6int | 0.1429 | ns | 0.17148 | ns |
|  |  | Aged vs Young | Aged GC-Bcl6no/low vs Young GC-Bcl6no/low | 0.5575 | ns | 0.5575 | ns |
|  |  |  | Aged GC-Bcl6int vs Young GC-Bcl6int | 0.0357 | * | 0.0714 | ns |
| Figure 2D | PD1hi - count per (µm2) | Aged | GC-Bcl6no/low vs GC-Bcl6high | 0.0002 | *** | 0.0009 | *** |
|  |  |  | GC-Bcl6no/low vs GC-Bcl6int | 0.0003 | *** | 0.0009 | *** |
|  |  |  | GC-Bcl6int vs GC-Bcl6high | 0.4286 | ns | 0.42886 | ns |
|  |  | Young | GC-Bcl6no/low vs GC-Bcl6int | 0.0714 | ns | 0.1428 | ns |
|  |  | Aged vs Young | Aged GC-Bcl6no/low vs Young GC-Bcl6no/low | 0.233 | ns | 0.3495 | ns |
|  |  |  | Aged GC-Bcl6int vs Young GC-Bcl6int | 0.3929 | ns | 0.42886 | ns |
|  | PD1hiKi67hi - count per (µm2) | Aged | GC-Bcl6no/low vs GC-Bcl6high | 0.0001 | *** | 0.0006 | *** |
|  |  |  | GC-Bcl6no/low vs GC-Bcl6int | 0.0011 | ** | 0.0033 | ** |
|  |  |  | GC-Bcl6int vs GC-Bcl6high | 0.9307 | ns | 0.9307 | ns |
|  |  | Young | GC-Bcl6no/low vs GC-Bcl6int | 0.0357 | * | 0.0714 | ns |
|  |  | Aged vs Young | Aged GC-Bcl6no/low vs Young GC-Bcl6no/low | 0.171 | ns | 0.2565 | ns |
|  |  |  | Aged GC-Bcl6int vs Young GC-Bcl6int | 0.25 | ns | 0.3 | ns |
|  | PD1hiCD57hi - count per (µm2) | Aged | GC-Bcl6no/low vs GC-Bcl6high | 0.0002 | *** | 0.0012 | *** |
|  |  |  | GC-Bcl6no/low vs GC-Bcl6int | 0.0123 | * | 0.0369 | * |
|  |  |  | GC-Bcl6int vs GC-Bcl6high | 0.9307 | ns | 0.9307 | ns |
|  |  | Young | GC-Bcl6no/low vs GC-Bcl6int | 0.0714 | ns | 0.1428 | ns |
|  |  | Aged vs Young | Aged GC-Bcl6no/low vs Young GC-Bcl6no/low | 0.1816 | ns | 0.2724 | ns |
|  |  |  | Aged GC-Bcl6int vs Young GC-Bcl6int | 0.3929 | ns | 0.47148 | ns |
|  | PD1hiKi67hiBcl6hi - count per (µm2) | Aged | GC-Bcl6no/low vs GC-Bcl6high | 0.0001 | *** | 0.0006 | *** |
|  |  |  | GC-Bcl6no/low vs GC-Bcl6int | 0.042 | * | 0.0792 | ns |
|  |  |  | GC-Bcl6int vs GC-Bcl6high | 0.7922 | ns | 0.7922 | ns |
|  |  | Young | GC-Bcl6no/low vs GC-Bcl6int | 0.0179 | * | 0.0537 | * |
|  |  | Aged vs Young | Aged GC-Bcl6no/low vs Young GC-Bcl6no/low | 0.0528 | ns | 0.0792 | ns |
|  |  |  | Aged GC-Bcl6int vs Young GC-Bcl6int | 0.3929 | ns | 0.47148 | ns |
| Figure 3D | Area aboce and below the curve (µm2)<br>GC B cells CD20hiKi67hiBcl6hi | x | Tonsils vs GC-Bcl6int | 0.0009 | *** | 0.00135 | *** |
|  |  |  | Tonsils vs GC-Bcl6high | 0.0004 | *** | 0.0012 | *** |
|  |  |  | GC-Bcl6int vs GC-Bcl6high | 0.8793 | ns | 0.8793 | ns |
|  | Area aboce and below the curve (µm2)<br>TFH cells PD1hi CD57hi | x | Tonsils vs GC-Bcl6int | 0.1475 | ns | 0.22125 | ns |
|  |  |  | Tonsils vs GC-Bcl6high | 0.0042 | ** | 0.0126 | ** |
|  |  |  | GC-Bcl6int vs GC-Bcl6high | 0.3434 | ns | 0.3434 | ns |
| Figure 5C | CD8hi Total cell counts | Aged | GC-Bcl6no/low vs GC-Bcl6high | 0.2129 | ns | 0.7404 | ns |
|  |  |  | GC-Bcl6no/low vs GC-Bcl6int | 0.8788 | ns | 0.9999 | ns |
|  |  |  | GC-Bcl6int vs GC-Bcl6high | 0.2468 | ns | 0.7404 | ns |
|  |  | Young | GC-Bcl6no/low vs GC-Bcl6int | >0.9999 | ns | 0.9999 | ns |
|  |  | Aged vs Young | Aged GC-Bcl6no/low vs Young GC-Bcl6no/low | 0.7214 | ns | 0.9999 | ns |
|  |  |  | Aged GC-Bcl6int vs Young GC-Bcl6int | 0.7857 | ns | 0.9999 | ns |
|  | CD8hi GrzBhi Total cell counts | Aged | GC-Bcl6no/low vs GC-Bcl6high | 0.0831 | ns | 0.4287 | ns |
|  |  |  | GC-Bcl6no/low vs GC-Bcl6int | 0.799 | ns | 0.799 | ns |
|  |  |  | GC-Bcl6int vs GC-Bcl6high | 0.4286 | ns | 0.799 | ns |
|  |  | Young | GC-Bcl6no/low vs GC-Bcl6int | 0.1429 | ns | 0.4287 | ns |
|  |  | Aged vs Young | Aged GC-Bcl6no/low vs Young GC-Bcl6no/low | 0.5743 | ns | 0.799 | ns |
|  |  |  | Aged GC-Bcl6int vs Young GC-Bcl6int | 0.7857 | ns | 0.799 | ns |
|  | MPOhi Total cell counts | Aged | GC-Bcl6no/low vs GC-Bcl6high | 0.5532 | ns | 0.66384 | ns |
|  |  |  | GC-Bcl6no/low vs GC-Bcl6int | 0.2786 | ns | 0.4179 | ns |
|  |  |  | GC-Bcl6int vs GC-Bcl6high | 0.0173 | * | 0.1038 | ns |
|  |  | Young | GC-Bcl6no/low vs GC-Bcl6int | 0.7857 | ns | 0.7857 | ns |
|  |  | Aged vs Young | Aged GC-Bcl6no/low vs Young GC-Bcl6no/low | 0.1946 | ns | 0.3892 | ns |
|  |  |  | Aged GC-Bcl6int vs Young GC-Bcl6int | 0.0357 | * | 0.1071 | ns |
|  | CD14hi Total cell counts | Aged | GC-Bcl6no/low vs GC-Bcl6high | 0.1246 | ns | 0.2492 | ns |
|  |  |  | GC-Bcl6no/low vs GC-Bcl6int | 0.799 | ns | 0.799 | ns |
|  |  |  | GC-Bcl6int vs GC-Bcl6high | 0.4286 | ns | 0.51432 | ns |
|  |  | Young | GC-Bcl6no/low vs GC-Bcl6int | 0.3929 | ns | 0.51432 | ns |
|  |  | Aged vs Young | Aged GC-Bcl6no/low vs Young GC-Bcl6no/low | 0.0365 | * | 0.1095 | ns |
|  |  |  | Aged GC-Bcl6int vs Young GC-Bcl6int | 0.0357 | * | 0.1095 | ns |
|  | CD68hi Total cell counts | Aged | GC-Bcl6no/low vs GC-Bcl6high | 0.0529 | ns | 0.07935 | ns |
|  |  |  | GC-Bcl6no/low vs GC-Bcl6int | 0.5058 | ns | 0.5058 | ns |
|  |  |  | GC-Bcl6int vs GC-Bcl6high | 0.1775 | ns | 0.213 | ns |
|  |  | Young | GC-Bcl6no/low vs GC-Bcl6int | 0.0357 | * | 0.0714 | ns |
|  |  | Aged vs Young | Aged GC-Bcl6no/low vs Young GC-Bcl6no/low | 0.0268 | * | 0.0714 | ns |
|  |  |  | Aged GC-Bcl6int vs Young GC-Bcl6int | 0.0357 | * | 0.0714 | ns |

**Table 7.**
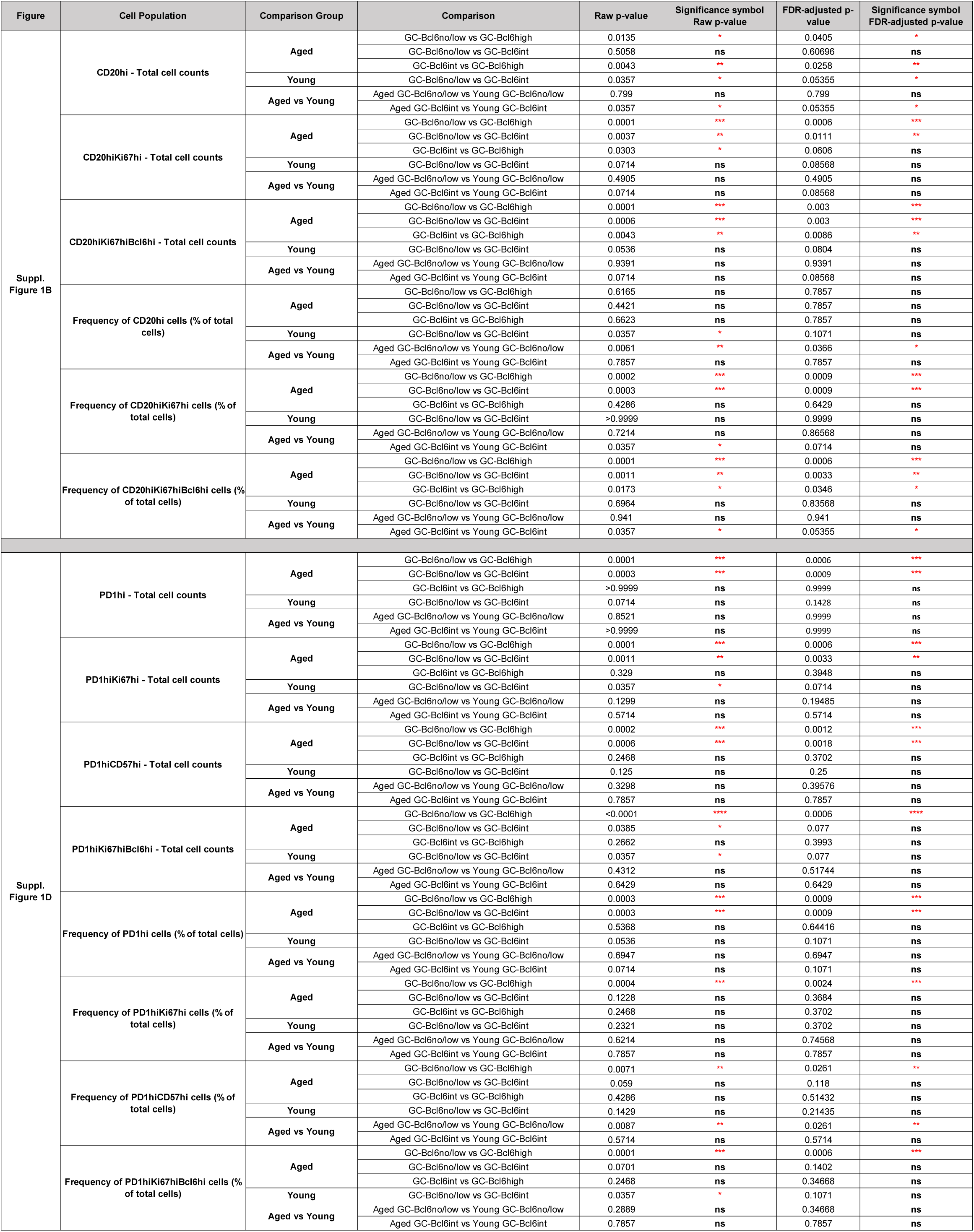
Summary of Statistical Analysis for Cell Population Comparisons: Raw and FDR-adjusted P-values with Significance Indicators Across Multiple Supplementary Figures.

## RESULTS

### COVID-19 infection induces diverse and heterogeneous RF profiling in LD-LNs

We started our investigation by analyzing F/GC B-cell subsets in LD-LNs from “Aged” (≥60 years) and “Young” (<57 years) individuals **(Tables 1 and 2)**, by applying a multiplex imaging assay **(Tables 3 and 4)** that allows for the simultaneous detection of CD20, Bcl6, and Ki67 **(Figures 1A and S1A)**. Follicular areas, identified by the density of CD20^high/dim^ cells, were further analyzed for B cell subsets using Histocytometry analysis for all combined follicles (total follicular area-All F) **(Figure 1B)**. Through visual assessment of raw mIF images and quantitative analysis of the Bcl6^high^ B cell counts, either per follicular area or total follicular area per tissue, we identified three distinct tissue subgroups: RF-Bcl6no/low, RF-Bcl6int, and RF-Bcl6high **(Figures 1C)**. Notably, the RF-Bcl6no/low group displayed Ki67 expression, indicating active proliferation and confirming these as RFs rather than primary or resting follicles. Furthermore, the area of individual RFs was significantly smaller in RF-Bcl6no/low tissues compared to those in the other two groups **(Figure 1D)**, while no significant differences were observed in other morphological features analyzed.

**Figure 1.**
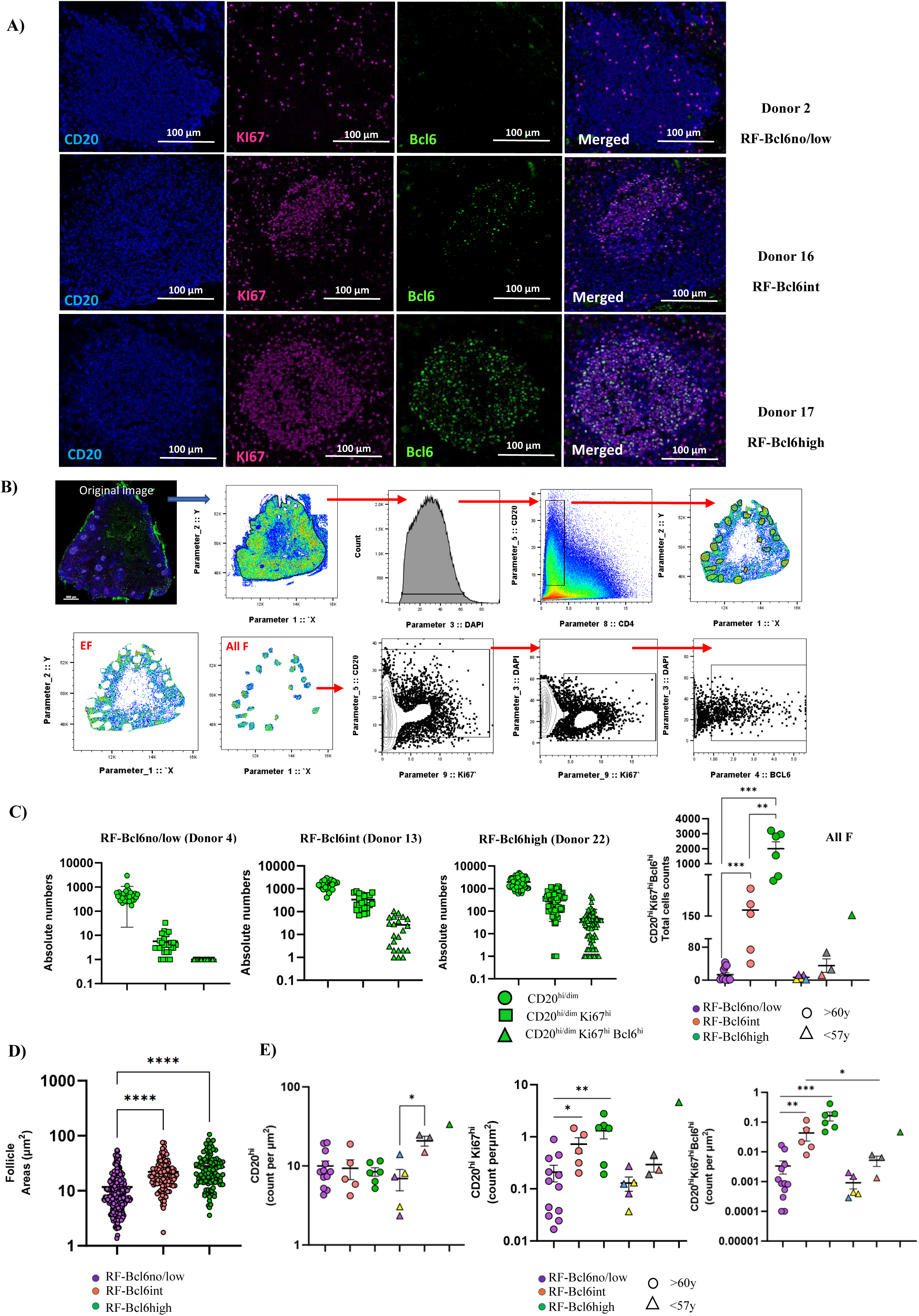
COVID-19 infection induces diverse and heterogeneous reactive follicles in LD-LNs. **A)** Representative fluorescence images (100µm) showing a follicular area in a RF-Bcl6no/low tissue (top images, donor 2), RF-Bcl6int tissue (middle images, donor 16), and in a RF-Bcl6high tissue (bottom images, donor 17) (CD20^high^-blue, Ki67^high^-magenta, Bcl6^high^-green, and a merged image). **B)** The gating scheme (Histocytometry) for the identification of B cell subsets is shown. Follicular areas were identified based on the density of CD20^high/dim^. The prevalence of B-cell subsets was calculated for all follicles combined (All F). **C)** Dot plot graphs showing the counted absolute numbers of B cell subsets (CD20^high/dim^, CD20^high/dim^Ki67^high^, CD20^high/dim^Ki67^high^Bcl6^high^) in three different donors (Donor 4 RF-Bcl6no/low, Donor 13 RF-Bcl6int and Donor 22 RF-Bcl6high), per follicle as well as for the total follicular area (All F, right panel). **D)** Dot plot graph of the calculated area of individual follicles normalized per tissue for RF-Bcl6no/low (purple), RF-Bcl6int (pink), and RF-Bcl6high (green) groups. **E)** Dot plot graphs showing the cell densities (normalized per µm^2^ cell counts) of B-cell subsets (CD20^high/dim^, CD20^high/dim^Ki67^high^, and CD20^high/dim^Ki67^high^Bcl6^high^) in the three groups for the “Aged” (round marks) and “Young” (triangle marks) individuals. Within the “Young” RF-Bcl6no/low group; the blue triangle depicts an axillary Lymph node (donor 25), and yellow triangles depict para-aortic and mediastinal lymph nodes from the same donor (donor 29). Within the “Young” RF-Bcl6int group, two different Mediastinal lymph nodes from the same donor (donor 24-a and donor 24-b) are depicted by the grey triangles. Asterisks denote p-value: * P≤0.05, ** P ≤ 0.01, and *** P ≤ 0.001 (Unpaired Non-Parametric Mann-Whitney T-Test).

Next, the cell densities (normalized cell counts/per µm2) of specific B-cell subsets were calculated. Despite the similar cell density of bulk CD20^high/dim^ B-cells across the three “Aged” subgroups, RF-Bcl6int and RF-Bcl6high tissues were characterized by significantly higher cell densities of proliferating Ki67^high^ and CD20^high/dim^Ki67^high^Bcl6^high^ B-cells **(Figure 1E)**. A similar profile was found when the frequency (%) or absolute cell counts of B-cell subsets **(Figure S1B)**. For the “Young” group, only a comparison between RF-Bcl6no/low and RF-Bcl6int subgroups could be applied. Although in “Aged” a significant difference was observed between RF-Bcl6no/low and RF-Bcl6int tissues in terms of bulk B-cell density, this difference did not reach significance when B-cell subsets were analyzed **(Figure 1E)**. A similar profile was observed when the frequency (% of total cells) or absolute cell counts of B-cell subsets were calculated **(Figure S1B)**.

Then, the pattern of the FDC network, as an additional surrogate of mature RFs, was also evaluated **(Figure S1C)**. We found a diverse expression of FDC in most RFs from both “Aged” RF-Bcl6no/low and RF-Bcl6high tissues. However, the FDC staining pattern varied between the two subgroups, with a connected network pattern more frequently observed in RF-Bcl6high tissues, while unconnected FDC positive events, a sign of less active follicles and mature RFs, were more commonly seen in RF-Bcl6no/low tissues **(Figure S1C)**. Altogether, our data suggest that RFs, as indicated by the cell densities of Ki67^high^ and Bcl6^high^ B-cells, in LD-LNs from COVID-19-infected individuals are highly diverse and heterogeneous.

### RF-Bcl6high LD-LNs harbor higher cell densities of proliferating Bcl6^high^ and CD57^high^ TFH cells

Given the critical role of TFH-cells in the development and maintenance of GC-RFs(5), we aimed to analyze their *in-situ* cell densities in the three aforementioned subgroups. TFH-cell subsets were identified based on the expression of PD1, Bcl6, Ki67, and CD57, a marker that marks a distinct TFH-cell subset(25), (26) **(Figures 2A and B)**. We noticed an inconsistency in the co-expression of CD57 and CD4 in the RFs among individuals, possibly reflecting a downregulation of CD4 receptor, similar to CD3 expression pattern(25), in these highly differentiated TFH cells. Given this characteristic, we refined our gating strategy to avoid bias by identifying TFH-cells as PD1^high^ and PD1^high^CD57^high^ **(Figure 2B)**. To set relevant cut-off values, we used PD1 and CD57 expression levels in extra-follicular (EF) CD4 T-cells as a reference and applied these thresholds to total F cells **(Figure 2B)**.

**Figure 2.**
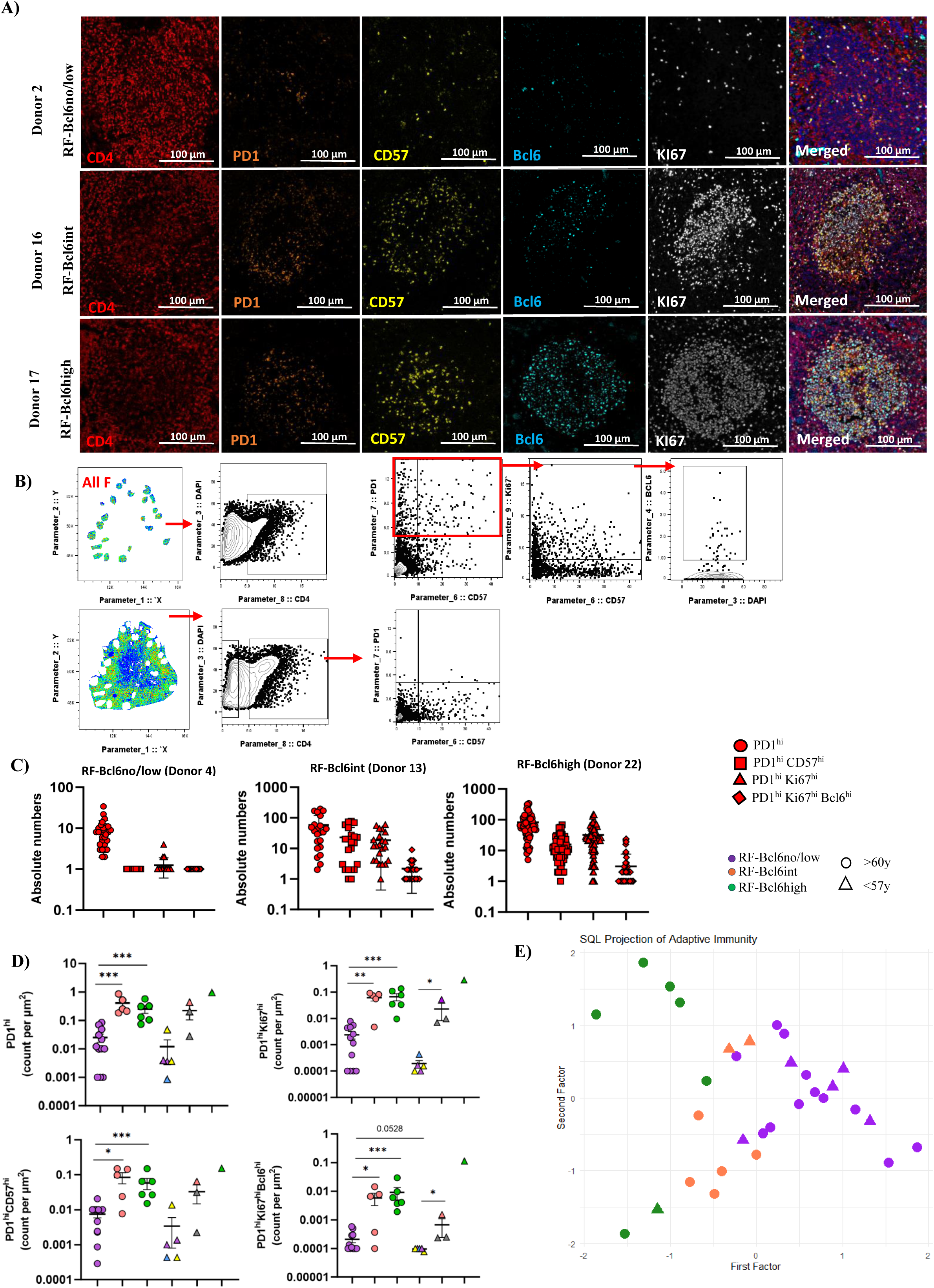
RF-Bcl6high LD-LNs harbor higher cell densities of proliferating Bcl6^high^ and CD57^high^ TFH cells. **A)** Representative fluorescence image (100 µm) showing a follicular area in a RF-Bcl6no/low tissue (left images, donor 2), RF-Bcl6int (middle image, donor 16), and a RF-Bcl6high tissue (right images, donor 17) (CD4^high^-red, PD1^high^-orange, CD57^high^-yellow, Bcl6^high^-cyan, Ki67^high^-white, and a merged image). **B)** The gating scheme for the identification of CD4 T cell subsets is shown. Follicular areas were identified by the expression of CD20^high/dim^. The prevalence of T cell subsets was calculated for all follicles combined (All F). **C)** Dot plot graphs showing the counted absolute numbers of T cell subsets (PD1^high^, PD1^high^CD57^high^, PD1^high^Ki67^high^, PD1^high^Ki67^high^Bcl6^high^) in three different donors (Donor 4 RF-Bcl6no/low, Donor 13 RF-Bcl6int and Donor 22 RF-Bcl6high) per follicle. **D)** Dot plot graphs showing the normalized (per µm^2^) counts of T cell subsets (PD1^high^, PD1^high^Ki67^high^, PD1^high^CD57^high^ and PD1^high^Ki67^high^Bcl6^high^) in the three groups for the “Aged” (round marks) and “Young” (triangle marks) individuals. Within the “Young” RF-Bcl6no/low group; the blue triangle depicts an axillary Lymph node (donor 25), and yellow triangles depict para-aortic and mediastinal lymph nodes from the same donor (donor 29). Within the “Young” RF-Bcl6int group, two different Mediastinal lymph nodes from the same donor (donor 24-a and donor 24-b) are depicted by the grey triangles. Asterisks denote p-value: * P≤0.05, ** P ≤ 0.01, and *** P ≤ 0.001(Unpaired Non-Parametric Mann-Whitney T-Test). **E)** SQL method clustering showing the projection of the different B cells and T cells subsets (adaptive immunity panel) based on CD20^high/dim^, CD20^high/dim^Ki67^high^, CD20^high/dim^Ki67^high^Bcl6^high^, CD4^high^, PD1^high^, PD1^high^Ki67^high^, PD1^high^CD57^high^ and PD1^high^Ki67^high^Bcl6^high^ cell counts in the three groups (RF-Bcl6no/low in purple, RF-Bcl6int in pink and, RF-Bcl6high in green), for the “Aged” (round marks) and “Young” (triangle marks).

Calculation of absolute numbers of TFH-cells positive for Bcl6 in individual RFs revealed a similar profile to the one found for the B-cells (**Figure 2C**). A significantly higher cell density of PD1^high^, PD1^high^Ki67^high^, PD1^high^CD57^high^, and PD1^high^Ki67^high^Bcl6^high^ cells in RF-Bcl6int and RF-Bcl6high compared to RF-Bcl6no/low tissues, in the “Aged” group, was observed **(Figure 2D)**. Despite the lack of statistical significance, the “Young” group exhibited a similar profile, with RF-Bcl6int showing significantly higher cell density of PD1^high^Ki67^high^ and PD1^high^CD57^high^ compared to RF-Bcl6no/low **(Figure 2D)**. A similar profile was found when the frequency (%) or absolute cell counts of TFH-cell subsets were analyzed **(Figure S1D)**. Then, we applied the SQL method(24) for clustering analysis of the tissues based on the B-cell and TFH-cell measurements. The SQL projection revealed a distinct clustering pattern for RF-Bcl6high compared to RF-Bcl6no/low and RF-Bcl6int subgroups **(Figure 2E)**. Altogether, our data show that the prevalence of Bcl6 in LD-LN from COVID-19 autopsies is associated with distinct RF immune landscaping.

### The increasing prevalence of Bcl6 is associated with an altered spatial distribution of RF immune cell subsets

Next, we analyzed the spatial distribution of key immune cell subsets within RFs in the “Aged” group by applying relevant algorithms to the RF-Bcl6int and RF-Bcl6high subgroups. Although not significant, a positive correlation, between PD1^high^CD57^high^ TFH-cells and CD20^high/dim^Ki67^high^Bcl6^high^ B-cells was found in the RF-Bcl6high subgroup **(Figure 3A)**. Furthermore, the ratio of CD20^high/dim^Ki67^high^Bcl6^high^ to PD1^high^CD57^high^ cells was significantly higher in the RF-Bcl6high compared to the RF-Bcl6int subgroup **(Figure 3B)** suggesting a higher possibility for B/T cell interaction in the RF-Bcl6high subgroup. Then, the X, and Y coordinates of individual cells (CD20^high/dim^Ki67^high^, CD20^high/dim^Ki67^high^Bcl6^high^, PD1^high^CD57^low^ and PD1^high^CD57^high^ cells) were extracted, focusing only on RFs containing at least 20 cells of the corresponding phenotypes. Using G-Function analysis **(Figures 3C and S2A)**, we found a significantly less scattered distribution for CD20^high/dim^Ki67^high^ B cells in the RF-Bcl6high compared to RF-Bcl6int tissues **(Figure 3D, upper left panel)** while the PD1^high^ TFH-cells, from the same RFs areas, exhibited the opposite profile **(Figure 3D, upper right panel)**. Then, RF areas were analyzed considering their Bcl6 reactivity. CD20^high/dim^Ki67^high^Bcl6^high^ B-cells displayed a significantly less scattered/higher clustering profile in tonsils compared to RF-Bcl6int and RF-Bcl6high tissues **(Figure 3D, lower left panel)**. This was also the case for PD1^high^CD57^high^ TFH-cells between tonsils and RF-Bcl6high LNs **(Figure 3D, lower right panel)**. Distance matrix analysis **(Figure 3E)** revealed a similar mean minimum Euclidean distances between proliferating CD20^high/dim^Ki67^high^ B-cells and PD1^high^CD57^low^ TFH-cells **(Figure 3F, left panel)** and a clear trend (p= 0.0784) for shorter distance between CD20^high/dim^Ki67^high^ B-cells and PD1^high^CD57^high^ TFH-cells **(Figure 3F, middle panel)** in RF-Bcl6high compared to RF-Bcl6int subgroup. However, no difference was found between CD20^high/dim^Ki67^high^Bcl6^high^ and PD1^high^CD57^high^ TFH-cells **(Figure 3F, right panel)**. Altogether, our findings suggest that between the RF-Bcl6int to RF-Bcl6high stage, there is a notable shift in the spatial organization of key immune cell subsets, with increased clustering of proliferating CD20^high/dim^Ki67^high^ B-cells and presumably higher possibility for their interaction with PD1^high^CD57^high^ TFH-cells in the RF-Bcl6high “Aged” subgroup of LD-LNs.

**Figure 3.**
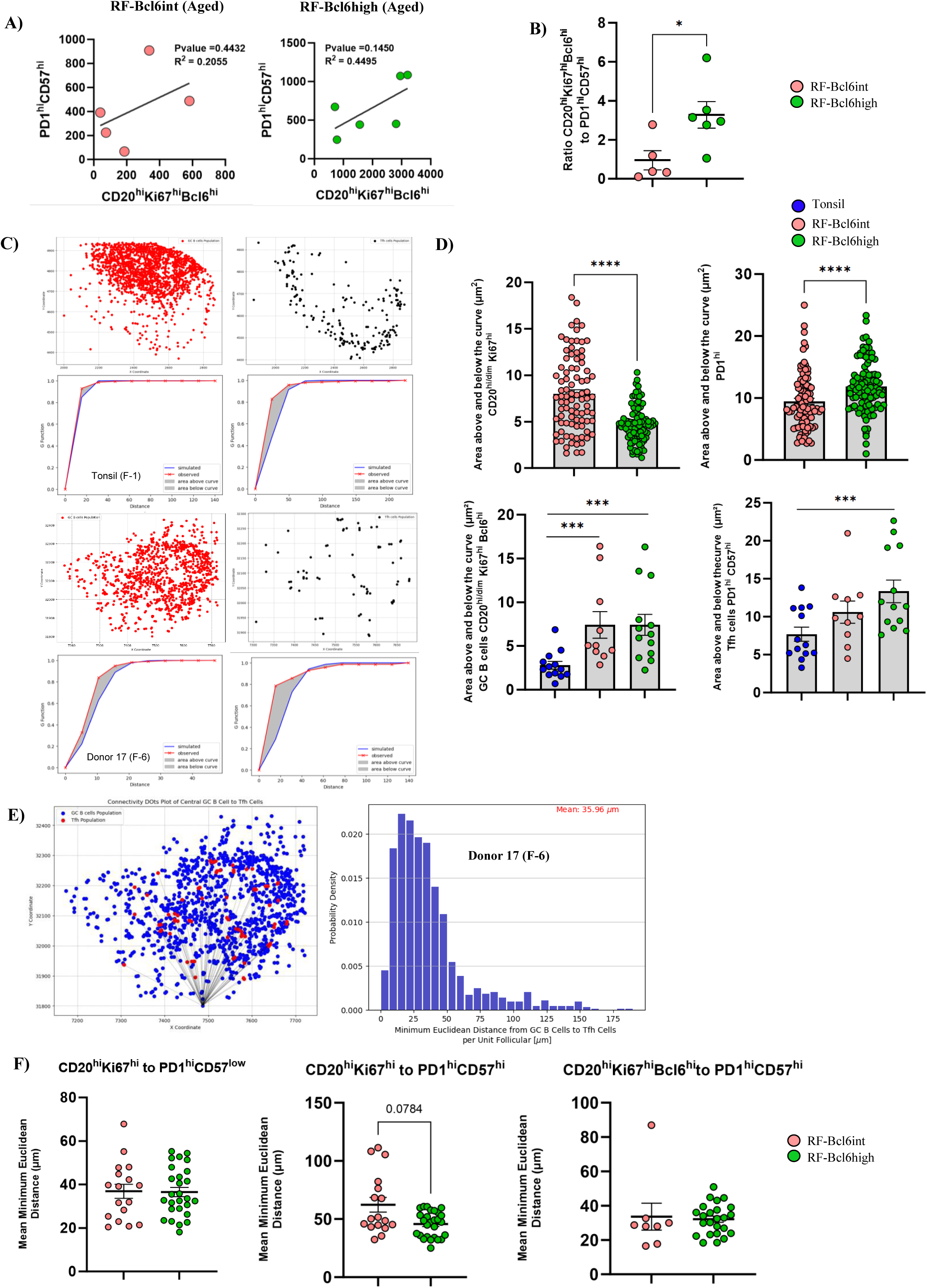
The increasing prevalence of Bcl6 is associated with an altered spatial distribution of RF immune cell subsets. **A)** Correlation analysis between PD1^high^CD57^high^ and CD20^high/dim^Ki67^high^Bcl6^high^ in “Aged” RF-Bcl6int (upper left), and “Aged” RF-Bcl6high (upper right) **B)** Ratio analysis of CD20^high/dim^Ki67^high^Bcl6^high^ to PD1^high^CD57^high^ cell counts in LD-LNs of “Aged” RF-Bcl6int (pink dots) and RF-Bcl6high (green dot); statistical analysis has been performed by Unpaired Non-Parametric Mann-Whitney T-Test. **C)** Graphs showing the distribution pattern of CD20^high/dim^Ki67^high^ (red) and PD1^high^CD57^high^ (black) cells in a control tonsil (top images, Tonsil F-1) and a RF-Bcl6high tissue (bottom images, Donor 17 F-6). Their corresponding G-Function graphs, showing the proximity of the G curve (red curve) to the theoretical curve (Poisson curve, blue curve), are shown too. The shade gray area represents the area between the two curves measured. **D)** Dot plot graphs of the measured area between the G and estimate curve for CD20^high/dim^Ki67^high^, PD1^high^, CD20^high/dim^Ki67^high^Bcl6^high^ and PD1^high^CD57^high^ for tonsils (blue dots), RF-Bcl6int (pink dots) and RF-Bcl6high (green dots) tissues. Asterisks denote p-value: * P≤0.05, ** P ≤ 0.01, and *** P ≤ 0.001 (Unpaired Non-Parametric Mann-Whitney T-Test). **E)** Connectivity dot plot identifying cell coordinates and distance measurements from CD20^high/dim^Ki67^high^Bcl6^high^ (blue dots) to PD1^high^CD57^high^ (red dots) cells in a follicular area of a RF-Bcl6high tissue (Donor 17 F-6). The corresponding bar graph shows the distribution of cells based on the distance from CD20^high/dim^Ki67^high^Bcl6^high^ to PD1^high^CD57^high^ cells. **F)** Dot plot graphs showing the mean of the minimum Euclidean distance measured from CD20^high/dim^Ki67^high^ to PD1^high^CD57^low^ or PD1^high^CD57^high^ and from CD20^high/dim^Ki67^high^Bcl6^high^ to PD1^high^CD57^high^ cells, in several follicular areas of RF-Bcl6int (pink dots) and RF-Bcl6high (green dots) tissues. Asterisks denote p-value: * P≤0.05, ** P ≤ 0.01, and *** P ≤ 0.001(Unpaired Non-Parametric Mann-Whitney T-Test).

### COVID-19 infection reveals a disconnection in RF immunoreactivity between LD-LNs and matched distal subdiaphragmatic LNs

We analyzed LD-LN and subdiaphragmatic (SD-LNs, serving as “control”) LNs from the same donors (**Table 5**) in the “Aged” group to investigate whether COVID-19 infection leads to a generalized RFs Bcl6 immunoreactivity. The adaptive immunity panel **(Table 3)** was applied to the SD-LNs **(Figure 4A)**. Overall, a trend for lower cell densities of B-cell and TFH-cell subsets that reached significance for the CD20^high/dim^Ki67^high^Bcl6^high^ cell subset in the SD-LNs compared to LD-LNs was found **(Figure 4B)**. Consistent with this profile, SD-LNs showed a significantly lower frequency of CD20^high/dim^Ki67^high^Bcl6^high^, PD1^high^Ki67^high^, and PD1^high^Ki67^high^Bcl6^high^ cells compared to LD-LNs **(Figure 4C)**. Therefore, the generalized inflammation associated with COVID-19 infection may not trigger the development of RF Bcl6 immune reactivity in distal, non-LD-LNs.

**Figure 4.**
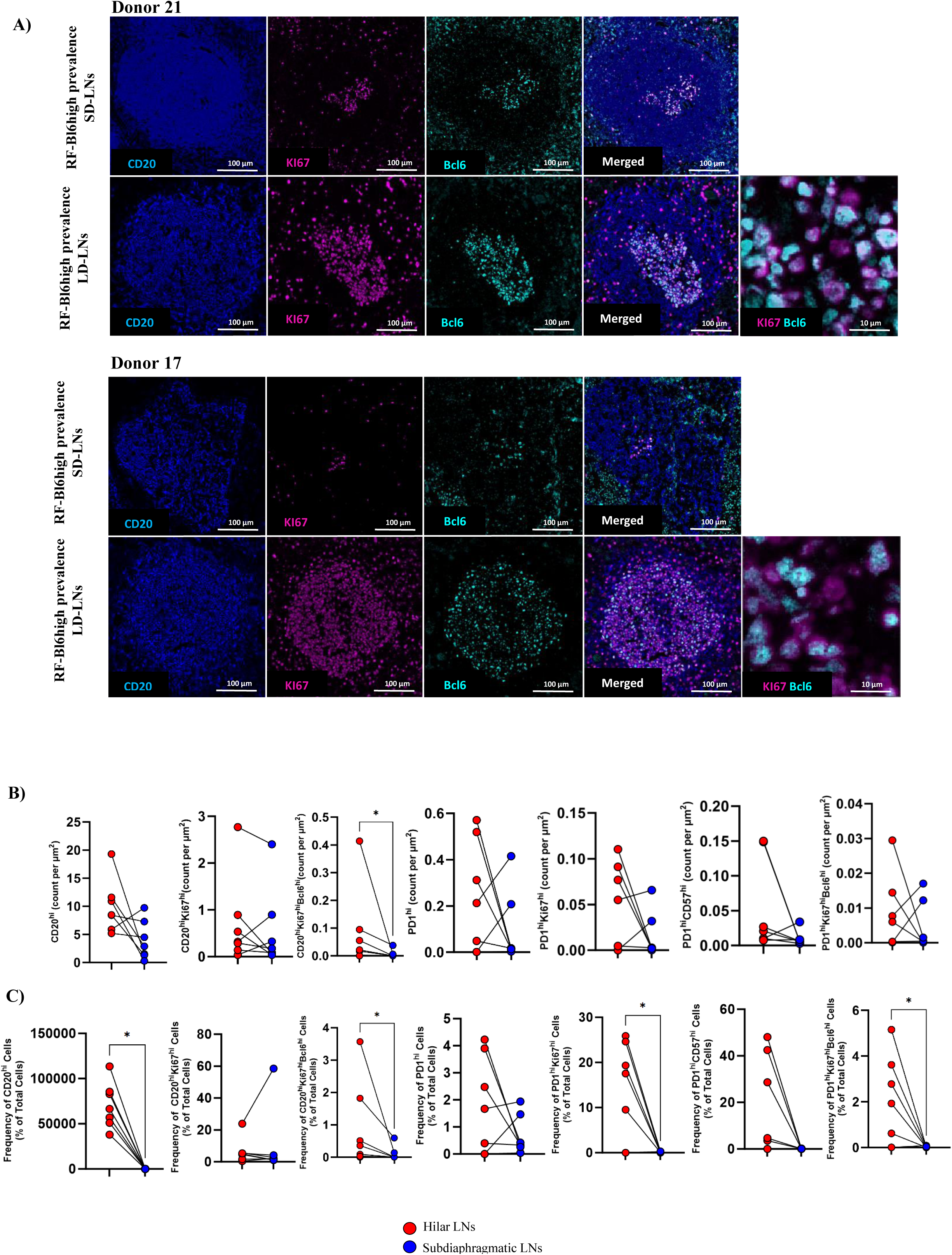
COVID-19 infection reveals a disconnection in F/GC immunoreactivity between LD-LNs and matched distal subdiaphragmatic LNs. **A)** Representative fluorescence images (100 µm and 10 µm) showing the CD20^high/dim^, Ki67^high^, and Bcl6^high^ expression in subdiaphragmatic and matched LD-LN follicular areas from two RF-Bcl6high samples (donor 21-upper panel and donor 17-lower panel) (CD20^high/dim^ -blue, Ki67^high^ -magenta, Bcl6^high^ - cyan, and a merged image). **B)** Dot plot graphs showing the cell densities normalized per µm^2^ cell counts of B and CD4 T cell subsets in the subdiaphragmatic (blue marks) and matched LD-LNs (red marks). Asterisks denote p-value: * P≤0.05, ** P ≤ 0.01, and *** P ≤ 0.001(Wilcoxon T-Test). **C)** Dot plot graphs showing the cell frequency (%) of total cells of B and CD4 T cell subsets in the subdiaphragmatic (blue marks) and matched LD-LNs (red marks). Asterisks denote p-value: * P≤0.05, ** P ≤ 0.01, and *** P ≤ 0.001(Wilcoxon T-Test).

### Contrary to B/TFH combined profile, LD-LNs RF-Bcl6high reactivity is not associated with a distinct CD8/innate immunity profile

Next, we investigated the prevalence of bulk and effector GrzB^high^ CD8^high^ T-cells, as well as innate immunity subsets (CD68, CD14 for monocytes/macrophages, and MPO, a marker of granulocytes) (**Figures 5A and S2B)**. The antibody panel **(Tables 3 and 4)** did not include follicular markers, so our Histocytometry analysis applied to both follicular (F) and extrafollicular (EF) areas **(Figure 5B)**. Similar cell densities of CD8^high^ and CD8^high^GrzB^high^ cells were found among the three tissue subgroups, for both “Young” and “Aged” individuals **(Figure 5C, left panels)**. In contrast to CD8^high^ T-cells, a mixed profile was observed when innate immunity cell types were analyzed. Significantly higher numbers of MPO^high^ and CD68^high^ cells were found in RF-Bcl6high compared to either RF-Bcl6int or RF-Bcl6no/low “Aged” donors (**Figure 5C, right panels)**. The comparison between matched subgroups showed significantly increased numbers of MPO^high^, CD68^high,^ and CD14^high^ cells in “Young” compared to “Aged” donors **(Figure 5C, right panels)**. Then, a SQL analysis based on the CD8, and innate immune cell subsets was applied. No distinct profile was found between the RF-Bcl6 subgroups, regardless of the aging **(Figure 5D)**. Thus, COVID-19 infection results in a relatively homogenous cell density profile of CD8/innate immunity cell types in LD-LNs regardless of their Bcl6 reactivity.

**Figure 5.**
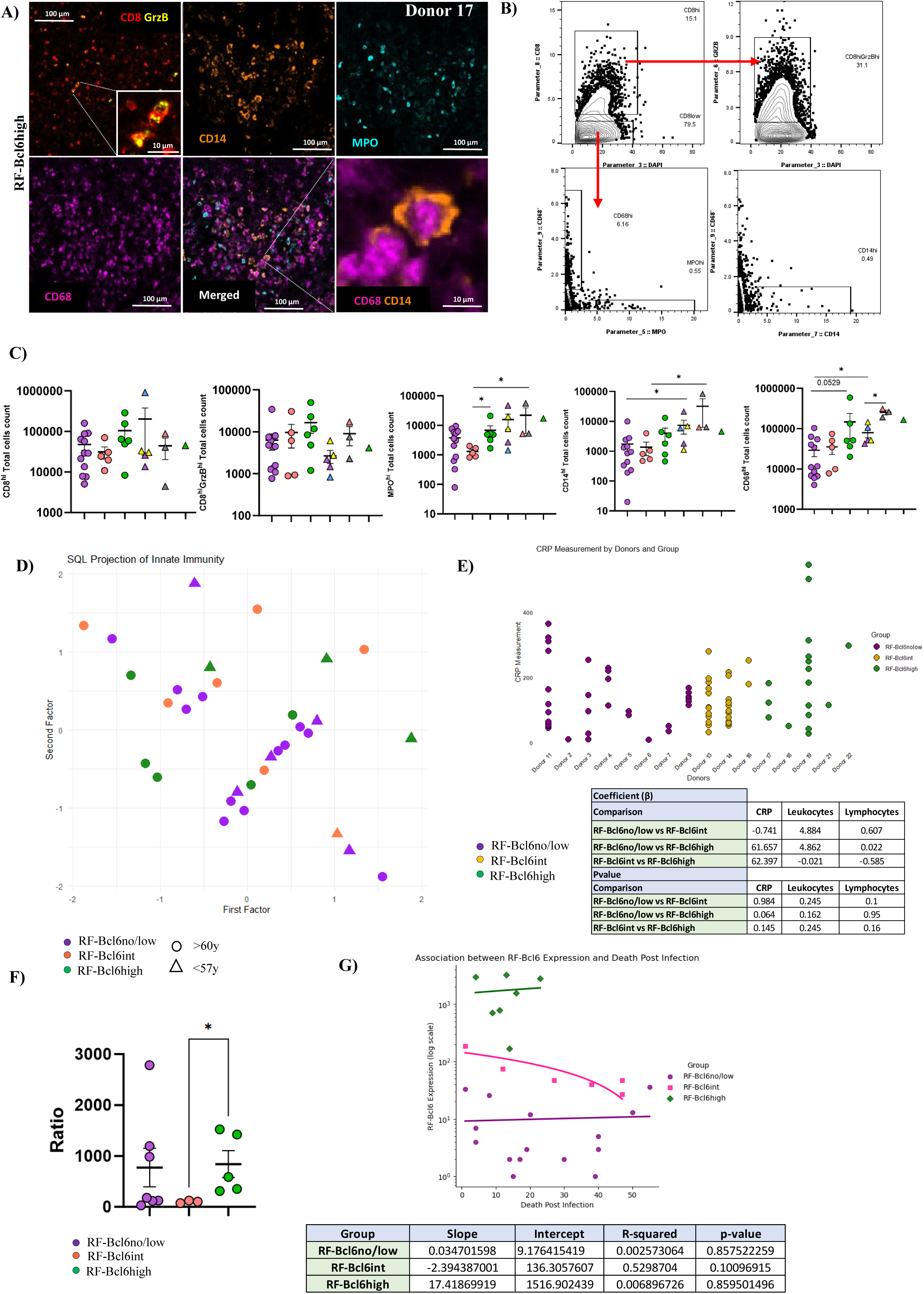
COVID-19 infection shows no distinct CD8 or innate immunity profile in RF-Bcl6high LD-LNs. **A)** Representative fluorescence images (100 µm and 10 µm) showing CD8^high^-red, GrzB^high^-yellow, CD14^high^-ceramide, MPO^high^-cyan, and CD68^high^-magenta expression in a follicular area from RF-Bcl6high tissue (Donor 17). Merged and zoomed areas are also shown. **B)** Gating scheme (Histocytometry) for identifying CD8^high^ T cells and innate immune cell subsets. **C)** Dot plots of total cell counts for CD8^high^, CD8^high^GrzB^high^, MPO^high^, CD14^high^, and CD68^high^ cells in the three groups for “Aged” (round) and “Young” (triangle) individuals. Within the “Young” RF-Bcl6no/low group, the blue triangle represents an axillary lymph node (donor 25), and yellow triangles represent para-aortic and mediastinal lymph nodes from donor 29. Within the “Young” RF-Bcl6int group, two different Mediastinal lymph nodes from the same donor (donor 24-a and donor 24-b) are depicted by the grey triangles. Asterisks denote p-values: * P≤0.05, ** P≤0.01, *** P≤0.001 (Mann-Whitney T-Test). **D)** SQL clustering projection of CD8^high^ T cells and innate immune cell subsets based on CD14^high^, CD68^high^, MPO^high^, CD8^high^, and CD8^high^GrzB^high^ counts cell counts in the three groups (RF-Bcl6no/low in purple, RF-Bcl6int in pink and, RF-Bcl6high in green), for the “Aged” (round marks) and “Young” (triangle marks). **E)** Dot plots showing CRP (mg/L) over time in “Aged” individuals from the three subgroups. Statistical analysis via Mixed Linear Regression, with summary in the lower panel. **F)** Ratio of LD-LNs MPO to circulating granulocytes (neutrophils, basophils, eosinophils) in the three subgroups. Asterisks denote p-values: * P≤0.05, ** P≤0.01, *** P≤0.001 (Mann-Whitney T-Test). **G)** Scatter plot with regression lines showing the association between RF-Bcl6 expression and death post-infection (DPI) across the three subgroups. RF-Bcl6 expression is on a logarithmic scale on the y-axis, and DPI on the x-axis. Data points: purple circles (RF-Bcl6no/low), pink squares (RF-Bcl6int), and green diamonds (RF-Bcl6high). Trend lines and a table summarizing key statistical parameters (slope, intercept, R-squared, p-value) are shown in the lower panel.

The possible association between serological measurement at different time points and RF-Bcl6 reactivity was investigated **(Figures 5E and S2C)**. To this end, a mixed linear regression model analysis was performed. A trend for higher levels of CRP was observed in RF-Bcl6high compared to RF-Bcl6no/low subgroup (β = 61.657, p = 0.064). No difference was found when leukocyte and lymphocyte counts were analyzed among the subgroups **(Figures 5E and S2C)**. Analysis of the ratio between LD-LN MPO^high^ cells and circulating granulocytes (neutrophils, basophils, eosinophils) showed a significantly higher ratio in RF-Bcl6high compared to RF-Bcl6int subgroup **(Figure 5F)**. Next, we sought to investigate whether RF-Bcl6 reactivity was (or not) associated with the estimated time after infection (based on days of hospitalization). No significant association was found in all three subgroups (p = 0.857, p = 0.1, and p = 0.859 for the RF-Bcl6no/low, RF-Bcl6int group, and RF-Bcl6high group respectively) **(Figure 5G)**. A similar profile was observed when the RF-Bcl6 expression over time was analyzed for “Aged” and “Young” donors separately **(Figure S2D)**. Our data suggest that the observed heterogeneity of LD-LN RF-Bcl6 reactivity may reflects the intrinsic capacity of the immune system to mount mature GC-RFs rather than a profile associated with the time of autopsy post-infection.

### The RF-Bcl6high tissues are characterized by a distinct *in situ* follicular transcriptomic profile

The GeoMx platform was applied to RF areas from tissues spanning the three subgroups, using normalized and batch-corrected data for analysis (**Figures S3A and S3B**). Principal Component Analysis (PCA) revealed distinct transcriptional profiles among RF areas based on Bcl6 expression **(Figure 6A)**. Given the tissue unavailability, we were not able to compare the *in situ* transcriptomic profiling between “Young” and “Aged” RF-Bcl6high groups. We focused our subsequent analysis on the “Aged” group by comparing RF-Bcl6no/low and RF-Bcl6int subgroups to the RF-Bcl6high one. Several differentially expressed genes (DEGs) emerged from the RF-Bcl6no/low vs. RF-Bcl6high comparison, with genes promoting GC-development (e.g. Bcl6, AICDA, IL21R, CXCL13, STAT3) overexpressed in RF-Bcl6high, while genes associated with a TH1-immune response (e.g. STAT4, TNFRSF10B, TNFRSF1A, CXCR3, TGFB3, IL7) were significantly upregulated in the RF-Bcl6no/low group **(Figures 6B and 6C)**. Analysis of the distribution “profile” of these gene sets among the RFs (ROIs) of a given tissue, showed a denser clustering for the “TH1-favoring” gene set in the RF-Bcl6no/low subgroup, compared to the more scattered profile for the “GC-favoring” gene set in the RF-Bcl6high subgroup **(Figures S3C and S4A)**. The comparison between RF-Bcl6int and RF-Bcl6high “Aged” subgroups revealed a significant upregulation of genes favoring TFH/GC formation (e.g. CXCL13, CD22, MIF, STAT3, AICDA) in the RF-Bcl6high group **(Figure 6D)**. The gradual upregulation of GC-promoting genes, such as AICDA, Bcl6, and STAT3, from RF-Bcl6no/low to RF-Bcl6int to RF-Bcl6high highlights the progressive development of maturing GC-RFs activity across these subgroups **(Figure 6E)**. Despite the consistent downregulation of genes associated with a TH1 response in the RF-Bcl6high compared to the RF-Bcl6no/low subgroup, a mixed profile was observed when the RF-Bcl6int subgroup was analyzed **(Figure 6E)**.

**Figure 6.**
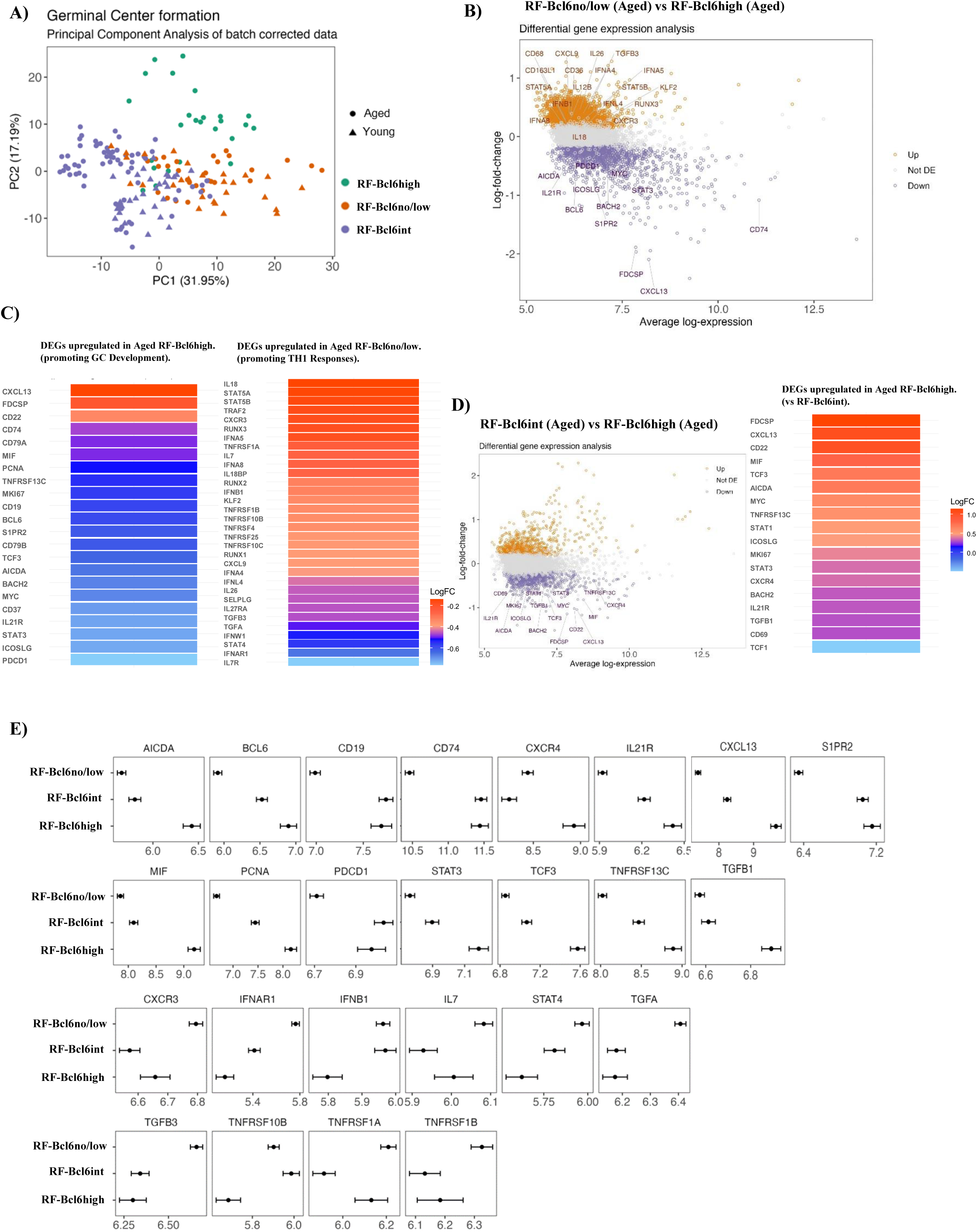
Distinct *in situ* transcriptomic profiles are associated with RF-Bcl6high follicles in LD-LNs. **A)** Principal component analysis (PCA) based on the 2 first components PC1 (31.95%) and PC2 (17.19%) of the gene expression after normalization and batch correction in the three groups, RF-Bcl6high (green dots), RF-Bcl6int (orange dots), and RF-Bcl6no/low (purple dots), for the “Aged” (round marks) and “Young” (triangle marks) individuals. **B)** Volcano plot showing differentially expressed genes in RF-Bcl6no/low vs RF-Bcl6high individuals from the “Aged” group. Orange dots present genes significantly increased in the RF-Bcl6no/low group and purple dots genes significantly increased in the RF-Bcl6high group. **C)** Heatmaps of differentially expressed genes (DEGs) promoting “GC development” upregulated in the RF-Bcl6high group (left image) or promoting “TH1 responses” upregulated in the RF-Bcl6no/low “Aged” group (right image). **D)** Volcano plot for DEGs between RF-Bcl6int and RF-Bcl6high individuals from the “Aged” group. Orange dots show genes significantly increased in the RF-Bcl6int group; while purple dots show genes significantly increased in the RF-Bcl6high of the “Aged” group. Heatmap of DEGs in RF-Bcl6high “Aged” individuals. **E)** Expression levels (log-transformed expression, x-axis) of specific genes across three RF-Bcl6 expression groups (y-axis).

Pathway enrichment analysis revealed upregulation of TNF-related pathways (e.g. “TNFR1 Signaling”, “NF-kappa-B Signaling”, “TRAIL Signaling”) and interferon pathways (e.g. “Alpha/Beta Interferon Signaling”) in follicles from “Aged” RF-Bcl6no/low compared to RF-Bcl6high tissues **(Figure 7A)**. Conversely, pathways related to metabolism, DNA repair, and immune activation (e.g. “Respiratory Electron Transport”, “ATP Synthesis”, “Mismatch Repair”) were significantly enriched in the RF-Bcl6high group. The “Aged” RF-Bcl6high subgroup also showed upregulation of RF-development pathways (e.g. “IL4”, “IL6 signaling”, “detoxification of reactive oxygen species”) compared to the “Aged” RF-Bcl6int subgroup **(Figure S4B)**.

**Figure 7.**
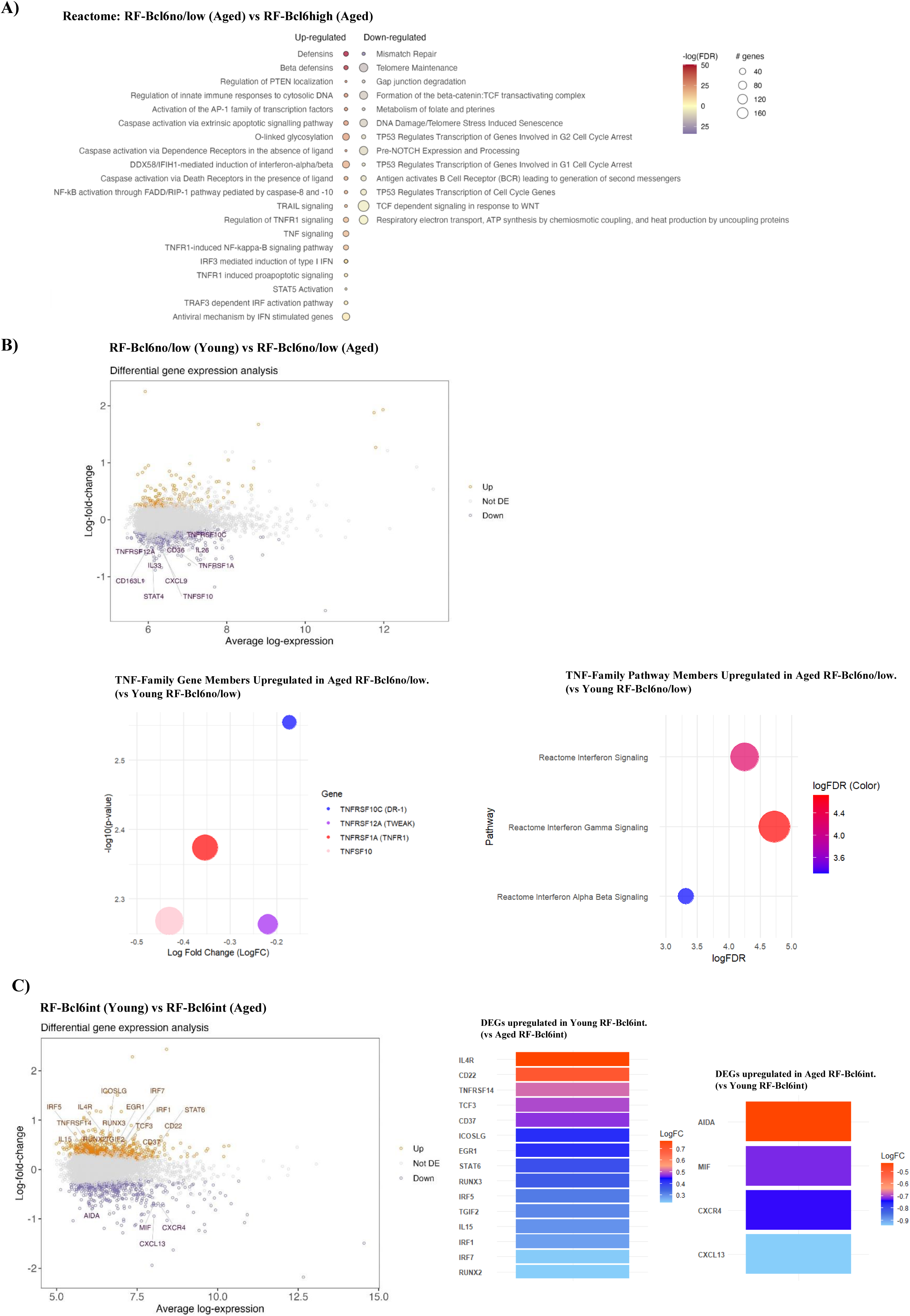
Altered follicular/GC transcriptomic profiling in LDLNs with respect to age. **A)** Dot plot of Reactome pathways significantly enriched in RF-Bcl6no/low vs RF-Bcl6high “Aged” individuals. The dot size represents the number of differentially expressed genes upregulated (left column) or downregulated (right column) in the RF-Bcl6no/low compared to RF-Bcl6high individuals. The color dots show the -log (FDR) representing the negative logarithm of the False Discovery Rate. **B)** Volcano plot for DEGs in RF-Bcl6no/low individuals from the “Young” and “Aged” subgroups. Orange dots show genes significantly increased in the “Young” RF-Bcl6no/low tissues and purple dots show genes significantly increased in the “Aged” RF-Bcl6no/low tissues. The bottom left panel displays a dot plot of TNF-family genes showing Log Fold Change (LogFC) and statistical significance (-log10(P-value)) in the “Aged” RF-Bcl6no/low group compared to the “Young” RF-Bcl6no/low group, with larger dots indicating higher significance and colors differentiating the genes. The bottom right panel features a bubble plot demonstrating the upregulation of “TNF-family-related” pathways, where the bubble size and position represent the degree of pathway activation and the color signifies FDR-adjusted significance levels.**C)** Volcano for DEGs in RF-Bcl6int individuals from the “Young” and “Aged” subgroups. The volcano plot is depicted with the log-fold change of each gene and the average log expression of each gene. Orange dots show genes highly expressed in “Young” RF-Bcl6int tissues and purple dots show genes highly expressed in “Aged” RF-Bcl6int tissues. Heatmap plots of DEGs highly expressed in the “Young” (left image) and the “Aged” (right image) RF-Bcl6int subgroup.

The differential expression analysis between “Young” and “Aged” RF-Bcl6no/low tissues revealed the upregulation of TNF-family genes (e.g. TNFSF10, TNFRSF1A, TNFRSF12A, TNFRSF10C) in the “Aged” group, indicating an enhanced pro-inflammatory environment **(Figure 7B)**. Pathway analysis supported this **(Figure 7B, left panel)**, further showing increased activation of interferon signaling pathways, crucial for driving inflammatory responses **(Figure 7B, right panel)**. Comparing “Young” and “Aged” RF-Bcl6int groups, the “Young” subgroup displayed upregulation of immune-related genes (e.g. IL4R, CD22, TNFRSF14), pointing to a more “reactive” immune environment, while the “Aged” subgroup exhibited higher expression of genes related to GC reactivity (AIDA, MIF, CXCL13, CXCR4) **(Figure 7C)**. Our “Young” RF-Bcl6no/low group included two different LD-LNs (24-a/24-b) from the same donor and anatomical site (mediastinal). No difference was observed in terms of DEGs or enriched pathways between these two LD-LNs (**Figure S4C**). Furthermore, the same group included one distal (axillary) LN, allowing for the comparison to the rest of the LD-LNs within the same (RF-Bcl6no/low) group **(Figure S4D)**. We observed several DEGs as well as an upregulation of pathways related to interleukin signaling (IL-7, IL-15, and IL17) and extracellular matrix organization in the axillary compared to LD-LNs **(Figure S4D)**. Therefore, the development (or not) of RF-Bcl6 reactivity in COVID-19 LD-LNs is associated with a distinct *in situ* molecular profiling that could be affected by aging.

“Aged” RF-Bcl6no/low tissues are characterized by a distinct *in situ* follicular macrophage profile.

Given the potential role of the aforementioned inflammatory pathways for the development of RFs in COVID-19, we aimed to further characterize the profile of macrophages in our tissue cohort. To this end, a set of genes from our GeoMx analysis was selected based on their macrophage “specificity” (including function-related genes like IL12B, CXCL9, and CCL7) and their expression was compared among different tissue subgroups. A significant upregulation of all selected genes was observed in the “Aged” RF-Bcl6no/low subgroup compared to the “Aged” RF-Bcl6high group **(Figure 8A)** while that was also true for a subset of the genes when a comparison to the “Young” RF-Bcl6no/low group was applied **(Figure 8B)**. Of note, we also found a significant (p=0.0069) association between the prevalence of CD68^high^ cells and death post-infection (DPI) only in the “Aged” RF-Bcl6no/low subgroup **(Figure 8C, left panel)**. A clear trend (p=0.066) was found for the CD14^high^ cell too **(Figure 8C, right panel)**. Therefore, our data indicate qualitative differences of the RF macrophages in the COVID-19 LD-LN with respect to Bcl6 reactivity.

**Figure 8.**
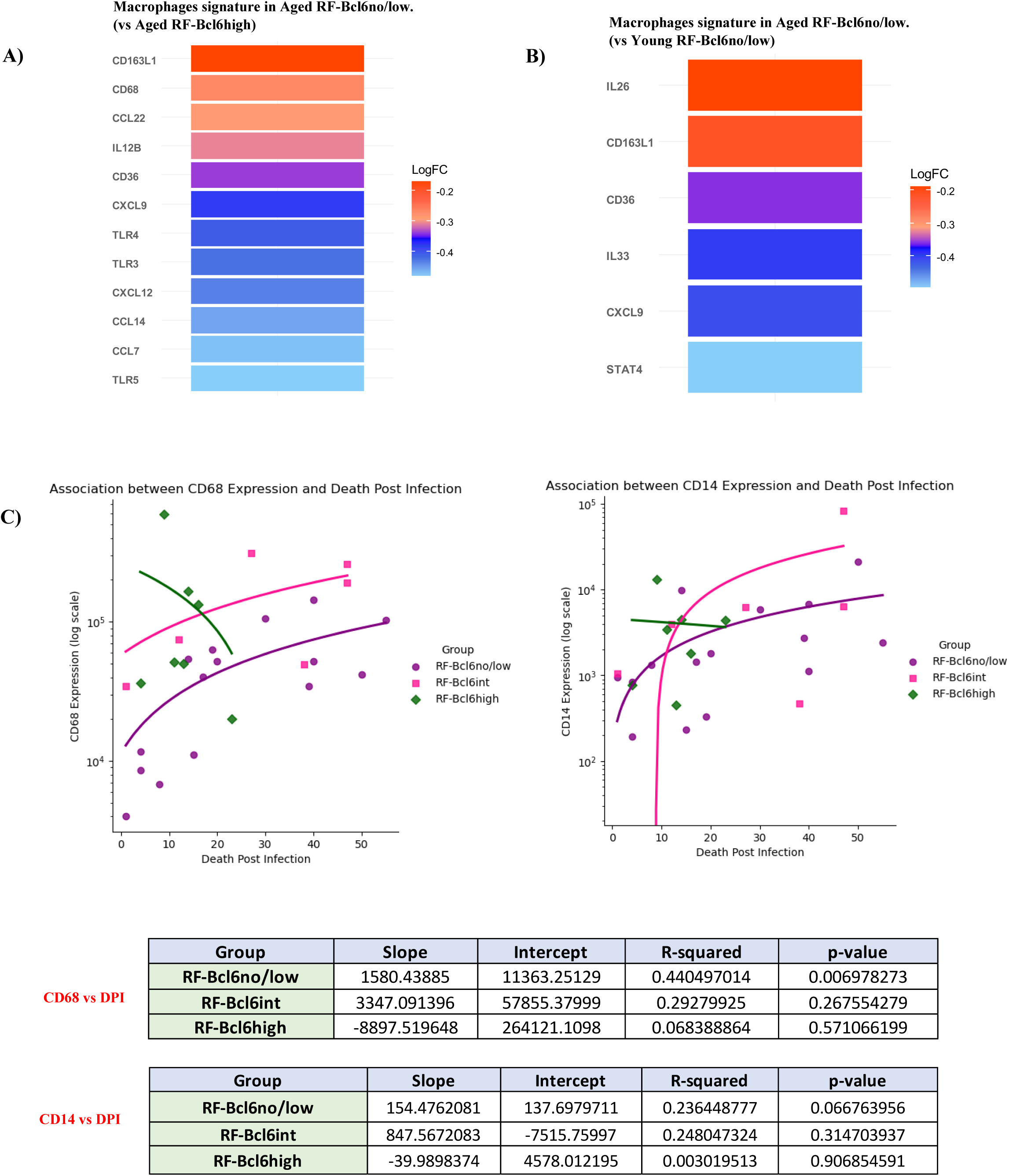
“Aged” RF-Bcl6no/low follicles exhibit a distinct *in situ* macrophage profile in COVID-19 LD-LNs. **A)** Heatmap illustrating macrophage-specific gene expression DEGs in RF-Bcl6no/low compared to RF-Bcl6high in LD-LNs from “Aged” COVID-19 infected individuals. **B)** Heatmap illustrating macrophage-specific gene expression DEGs in “Aged” compared to “Young” RF-Bcl6no/low in LD-LNs COVID-19 infected individuals. **C)** Linear regression analysis showing the association between the cell densities of CD68 and death post-infection (DPI) (left panel) or CD14 and DPI (right panel) in LD-LNs from “Aged” COVID-19-infected individuals. CD68 and CD14 cell densities are on a logarithmic scale on the y-axis, and DPI on the x-axis. Data points: purple circles (RF-Bcl6no/low), pink squares (RF-Bcl6int), and green diamonds (RF-Bcl6high). A table summarizing key statistical parameters for each group, including the slope, intercept, R-squared, and p-value based on the linear regression analysis is shown too (lower panel).

## Discussion

Given the lack of accessibility to relevant human LNs, the cellular and molecular mechanisms regulating the follicular/germinal center immune dynamics in draining LNs during viral infections, particularly during the early phase— the initial period following viral exposure when the immune response is first activated—are poorly understood. Here, we leveraged the availability of LD-LNs and matched control distal LNs from COVID-19 autopsies to investigate the RFs’ immune landscaping and the molecular pathways active *in situ* with a focus on the impact of donor aging. The “Young” group included only one RF-Bcl6high LD-LN, challenging the analysis between “Young” and “Aged” donors for this subgroup.

We stratified LD-LNs into three distinct subgroups based on Bcl6 expression levels to assess RF dynamics and activity. Analyzing the RFs based on CD20^high/dim^ expression, we detected Ki67 expression, indicating active proliferation and confirming these structures as secondary follicular areas rather than primary, resting follicles in all tissues analyzed. The three subgroups (RF-Bcl6no/low, RF-Bcl6int, and RF-Bcl6high) were defined by varying levels of Bcl6 expression. Analysis of individual RFs in the RF-Bcl6int and RF-Bcl6high subgroups revealed substantial variability in Bcl6 expression, even within the same tissue or donor. The RF-Bcl6no/low subgroup was characterized by i) significantly smaller RFs areas, ii) the presence of proliferating B-cells (CD20^high/dim^Ki67^high^), although at significantly lower levels than in the other two subgroups, and iii) a distinct FDC staining pattern(27) that further highlighted these structures as reactive, less matured rather than resting follicles. Additionally, we observed inconsistent co-expression of CD4 and CD57 in the RFs, likely due to CD4 downregulation in PD1^high^CD57^high^ TFH cells. Therefore, we chose to analyze total PD1^high^CD57^high^ and PD1^high^CD57^low^ cells within the RFs as a surrogate marker for TFH cells. The significantly higher PD1 expression in RFs versus extrafollicular areas, along with the significantly lower PD1 expression in extrafollicular and follicular CD8^high^ T-cells compared to TFH cells(28), suggests that our approach would not lead to miscalculation of TFH cells. Taking into consideration the calculated B and CD4 T-cell subsets, a distinct clustering of RF-Bcl6high tissues was observed. This clustering was less evident between RF-Bcl6no/low and RF-Bcl6int tissues, possibly reflecting an intermediate stage of germinal center development in the RF-Bcl6int subgroup. Overall, our data revealed a balanced representation of the three RF subgroups based on Bcl6 expression in COVID-19-infected individuals.

Next, we sought to investigate the spatial positioning of CD4 and B-cell subsets in RF-Bcl6int and RF-Bcl6high LD-LNs from “Aged” donors. Despite the similar cell densities of CD20^high/dim^Ki67^high^ and PD1^high^ cells among the two subgroups, their distribution/scattering profile was different. The PD1^high^CD57^high^ phenotype marks a unique TFH subset with a distinct function, positioning (closer to Dark Zone compared to PD1^high^CD57^low^ TFH cells (25)), and molecular profile(29), (30), (25). A clear trend for closer proximity of PD1^high^CD57^high^ TFH to CD20^high/dim^ Ki67^high^ B cells was observed in RF-Bcl6high tissues compared to RF-Bcl6int tissues. Assuming that the distance between two cells reflects the likelihood of their interaction, these findings suggest an increased possibility for B/T cell interactions in the RF-Bcl6high subgroup. The ratio and spatial positioning profile of B-cell and TFH-cell subsets further supports our hypothesis that the RF-Bcl6int phenotype may represent a transitional or less mature stage of RF development compared to the more established RF-Bcl6high phenotype.

The direct comparison between LD-LNs and matched subdiaphragmatic, distal LNs, suggests that despite the systemic inflammation/immune activation associated with COVID-19 and the possible dissemination of the virus at different anatomical sites across the human body(31), COVID-19 infection can induce mature RFs immune reactivity selectively in regional/draining lymphoid organs. The formation of Tertiary Lymphoid Structures (TLS) in long COVID-19 was recently described(32). Whether the development of a RF-Bcl6high reactivity in LD-LNs is also associated with the presence of lung-associated TLS, in our cohort, remains to be determined.

*In situ* analysis of bulk and effector (GrzB^high^) CD8 T-cells revealed a similar profile across the groups, irrespective of aging. Contrary to B-cells and T-cells, clustering analysis considering CD8, and innate cell types showed a more homogeneous profile among the RF-Bcl6 groups, likely reflecting a generalized inflammatory LN environment characteristic in COVID-19 infection. Among the serum measurements, CRP was found significantly higher in RF-Bcl6high compared to RF-Bcl6no/low tissues. This profile, however, was not associated with lymphopenia or higher circulating numbers of neutrophils, a hyperinflammation profile previously described in COVID-19(33). The higher ratio of LN MPO / circulating granulocytes found in RF-Bcl6high tissues may represent a higher extravasation of this cell type to LD-LNs. Most importantly, our modeling analysis revealed that the Bcl6 cell density in LD-LNs is not associated with the time of death post-infection (DPI) suggesting that the intrinsic ability of the immune system of a given individual and *in situ* operating biological factors are responsible for the observed phenotypes rather than the lack of adequate time for the development of RF-Bcl6int/high reactivity.

The spatial follicular transcriptomic profiling showed distinct molecular signatures associated with the level of RF-Bcl6 expression in “Aged” lymph node (LD-LN) tissues. Previous studies, using bulk LN mRNA preparation, have attributed the absence of Bcl6 expression in COVID-19 hilar LNs to overexpression of TNF-related genes, specifically TNF-α(6), which impaired the development of TFH cells. Our data extend these findings by showing that compromised Bcl6 expression in LD-LNs is associated with significantly increased expression of genes and pathways favoring TH1 differentiation within the follicular/GC areas. Specifically, genes counteracting TFH-development, such as KLF2(34), STAT4(35), STAT5A/B(36), IFNAR1, and TGFB3(37), as well as those characterizing or favoring a TH1-response, including CXCR3 and TNF-related genes (TNF-receptors like TNFRSF1A (DR1), TNFRSF25 (DR3), and TNFRSF10B (DR5), and TNF-signaling mediators like TRAF2). Additionally, other key genes involved in TNF-signaling, such as TNFRSF1B (TNFR2) and TNFRSF10C (DcR1), contribute to the regulation of immune responses and apoptosis pathways, were overexpressed in RF-Bcl6no/low compared to RF-Bcl6high tissues. Conversely, the RF-Bcl6high phenotype was supported by significant increases in genes favoring B/TFH cell development (e.g. BCL6, AICDA, CD74, STAT3), their trafficking (e.g. CXCL13, CXCR4, S1PR), T-cell development (TCF3), and pathways like “Antigen BCR activation” and “TCF-dependent signaling”. Therefore, our imaging data, coupled with transcriptomic analysis, suggest that high Bcl6 cell density is possibly associated with the coordinated expression and function of several cell subsets and molecular pathways favoring GC development. On the other hand, the RF-Bcl6int profile was characterized by an intermediate expression of genes favoring GC development and a mixed expression profile of TH1-favoring genes. These findings further support the hypothesis that RF-Bcl6int phenotype represents a transitional stage of GC development, in line with the aforementioned B-cell and T cell-based clustering profiles and neighboring analysis.

Our imaging data point to an overall trend for higher cell densities of macrophage/monocytes in RF-Bcl6high compared to RF-Bcl6no/low tissues. However, the expression pattern of macrophage function-related genes suggests an elevated capacity of macrophages for stimulation (expression of TLR3) and production of cytokines/chemokines, in the RF-Bcl6no/low subgroup. Furthermore, the modeling analysis revealed an accumulation of CD68 and CD14 over time DPI only in the RF-Bcl6no/low subgroup. Our data are in line with previous findings showing that elevated TNF-α production in COVID-19 LNs contributes to the loss of GC Bcl6 reactivity(6). Therefore, the quality of macrophages, the primary producers of TNF-α(38), could play a critical role in the development of GCs during early viral infections. Further, we observed upregulation of certain macrophage function-related genes in “Aged” RFs areas, likely in response to chronic low-grade inflammation, often referred to as "inflammaging”, which might reflect the heightened inflammatory environment in aging lymph nodes(39). These findings highlight the crucial role the macrophages could play in shaping the aging germinal center microenvironment and modulation of pathogen-specific B-cell responses.

We should also highlight the limitations of the current study including the absence of “non-infected control tissues” as well as the lack of data on antibody responses (titers or affinity). Whether the described RF-Bcl6 profile is linked to the development of anti-COVID-19 antibody responses, particularly those of high-affinity, cannot be addressed by the current study. However, our study provides cellular and molecular profiles, associated with the human F/GC development, that could fuel future investigation aiming to further understand the F/GC immune landscaping and model the development of B-cell responses in health and disease(40). This is of particular interest for the early phase of viral infections where accessibility to relevant LNs is highly challenging if possible.

---

**Suppl. Figure 1.**
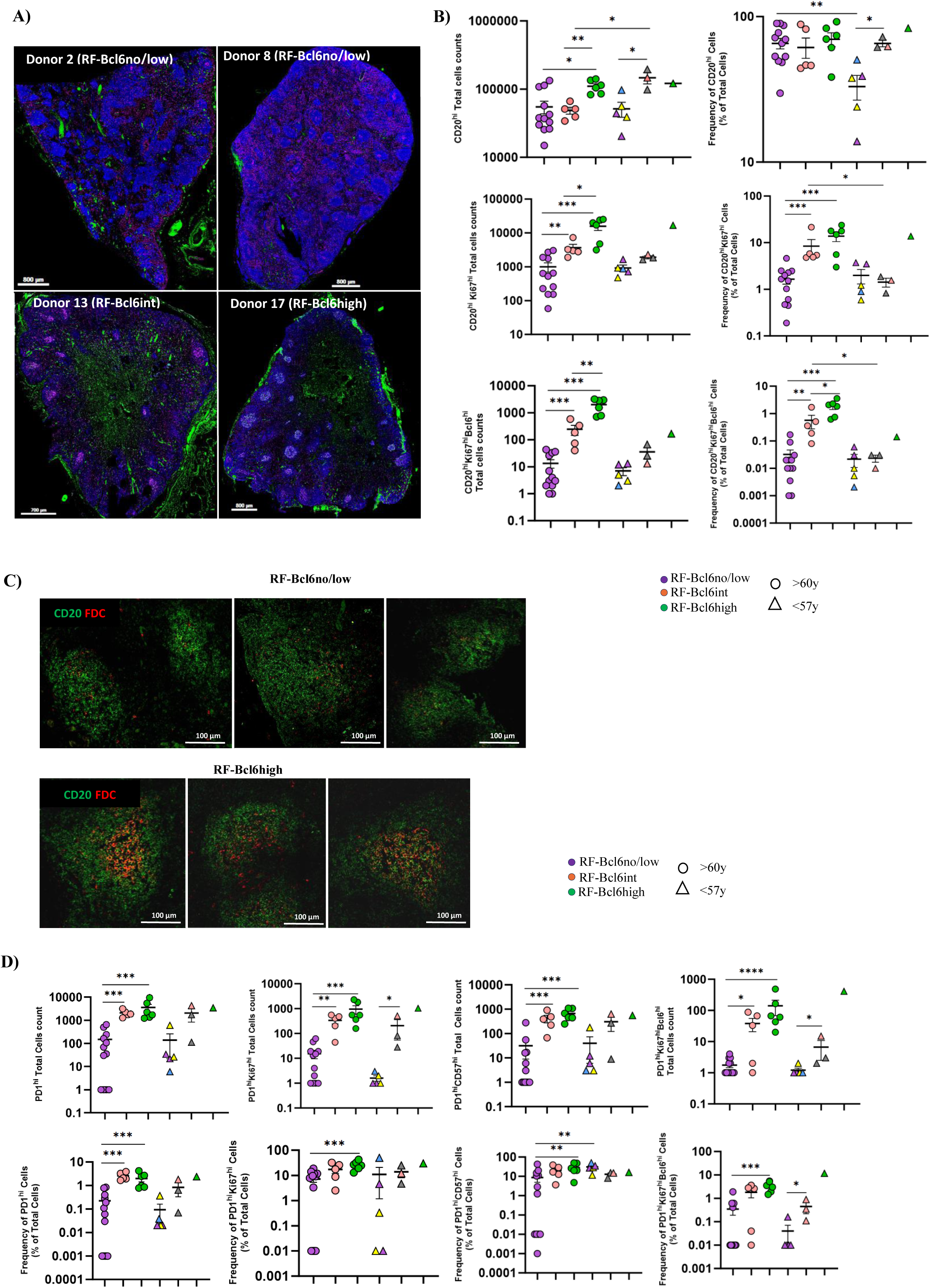
**A)** Representative fluorescence images showing tissues (overview) from each group in a RF-Bcl6no/low tissue (Donor 2 and Donor 8), RF-Bcl6int (Donor 13), and RF-Bcl6high (Donor 17). **B)** Dot plot graphs showing the total cell counts and frequency (%) of total cells of B cell subsets (CD20^high/dim^, CD20^high/dim^Ki67^high^, and CD20^high/dim^Ki6^high^Bcl6^high^) in the three groups for the “Aged” (round marks) and “Young” (triangle marks) individuals. Within the “Young” RF-Bcl6no/low group, the blue triangle represents an axillary lymph node (donor 25), and the yellow triangles represent para-aortic and mediastinal lymph nodes from donor 29. Within the “Young” RF-Bcl6int group, two different Mediastinal lymph nodes from the same donor (donor 24-a and donor 24-b) are depicted by the grey triangles. Asterisks denote p-values: * P≤0.05, ** P≤0.01, *** P≤0.001 (Mann-Whitney T-Test). **C)** Representative fluorescence images (100 µm) showing the expression of FDC (red) in follicular areas (CD20, green) from RF-Bcl6no/low (n=3) and RF-Bcl6high (n=3) tissues. **D)** Dot plot graphs showing the total cell counts and frequency (%) of total cells, of T cell subsets (PD1^high^, PD1^high^Ki67^high^, PD1^high^CD57^high^, and PD1^high^Ki67^high^Bcl6^high^) in the three groups for the “Aged” (round marks) and “Young” (triangle marks) individuals. Within the “Young” RF-Bcl6no/low group, the blue triangle represents an axillary lymph node (donor 25), and the yellow triangles represent para-aortic and mediastinal lymph nodes from donor 29. Within the “Young” RF-Bcl6int group, two different Mediastinal lymph nodes from the same donor (donor 24-a and donor 24-b) are depicted by the grey triangles. Asterisks denote p-values: * P≤0.05, ** P≤0.01, *** P≤0.001 (Mann-Whitney T-Test).

**Suppl. Figure 2.**
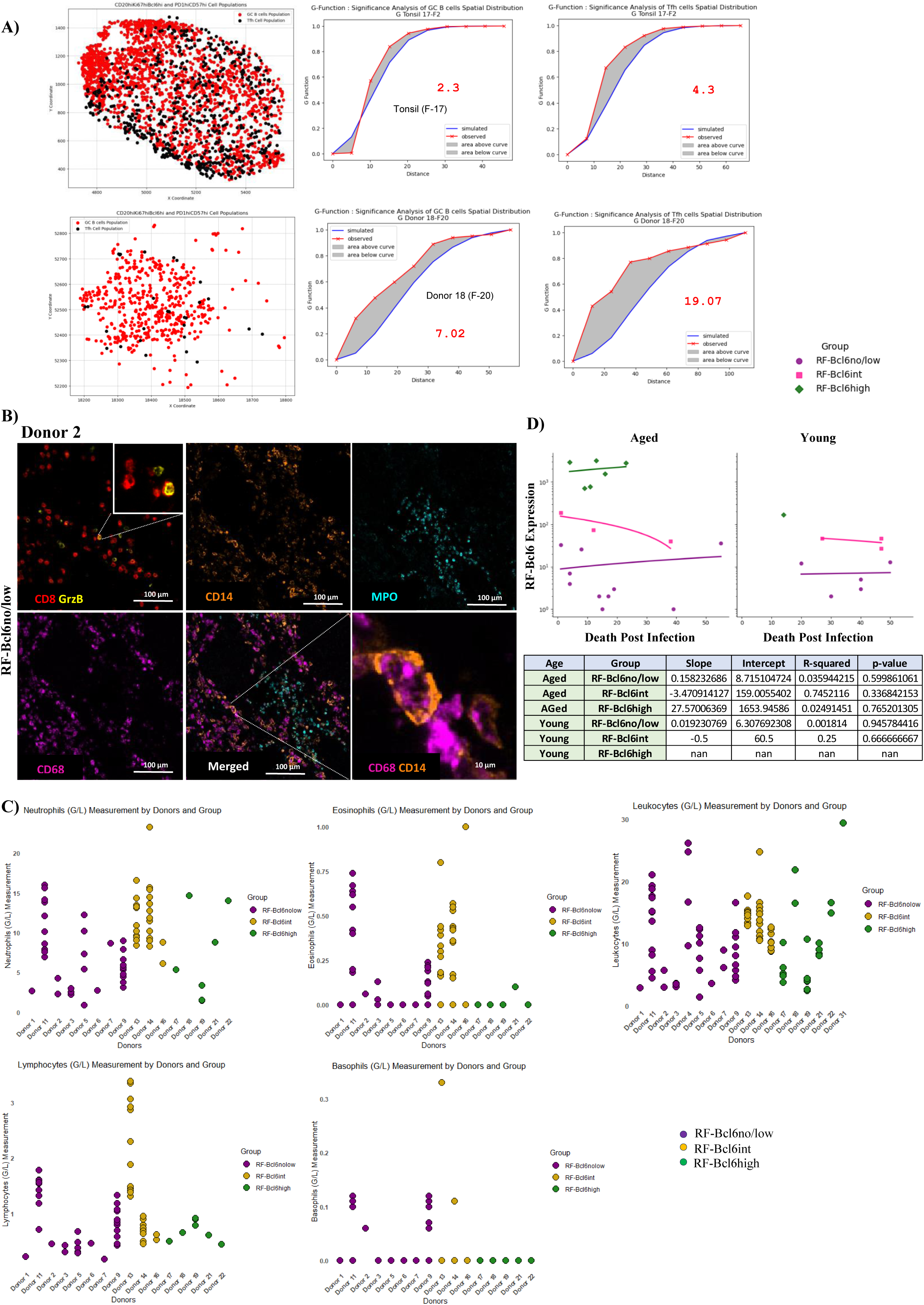
**A)** Point pattern distribution graphs showing a follicular area in control tonsillar tissue (top, Tonsil 17 – F2) and in a RF-Bcl6high tissue (bottom, Donor 18 – F20) (CD20^high/dim^Ki67^high^Bcl6^high^-red and PD1^high^CD57^high^-black) and their corresponding graphic representation of the G-Function analysis showing the proximity of the G curve (red curve) to the theoretical estimate curve (Poisson curve, blue curve). The shade gray area represents the area between the two curves measured. **B)** Representative fluorescence image (100µm) showing a follicular area in a RF-Bcl6no/low tissue (Donor 2); CD8^high^-red and GrzB^high^-yellow (insert: a zoomed area is shown, 10µm), CD14^high^-orange, MPO^high^-cyan, CD68^high^-magenta as well as a merged image are shown. The zoomed image shows the localization of CD68^high^-magenta and CD14^high^-orange. **C)** Dot plots showing serological measurement for total Neutrophils (G/L), Eosinophils (G/L), Leukocytes (G/L), Lymphocytes (G/L) and Basophils (G/L) at different time points in the group of “Aged” individuals for RF-Bcl6no/low (purple dots), RF-Bcl6int (yellow dots) and RF-Bcl6high (green dots). **D)** Scatter plot with regression lines illustrating the association between RF-Bcl6 expression and death post-infection (DPI) across the three subgroups from “Aged” (left panel) and “Young” (right panel). RF-Bcl6 expressions are plotted on a logarithmic scale on the y-axis, while DPI is shown on the x-axis. Data points for the RF-Bcl6no/low group are represented by purple circles, for the RF-Bcl6int group by pink squares, and for the RF-Bcl6high group by green diamonds. Summary table of the Mixed Linear Regression Model performed to compare RF-Bcl6no/low (reference) vs RF-Bcl6int, RF-Bcl6no/low (reference) vs RF-Bcl6high, and RF-Bcl6int (reference) vs RF-Bcl6high from the “Aged” group for the measurement of CRP. The results, including p-values and intercept (β) values, are summarized in the table. The intercept (β) represents the average outcome value for the reference group when all other predictors are zero. A p-value less than 0.05 indicates a statistically significant difference between the groups. Statistical analysis has been conducted using a Mixed Linear Regression model.

**Suppl. Figure 3.**
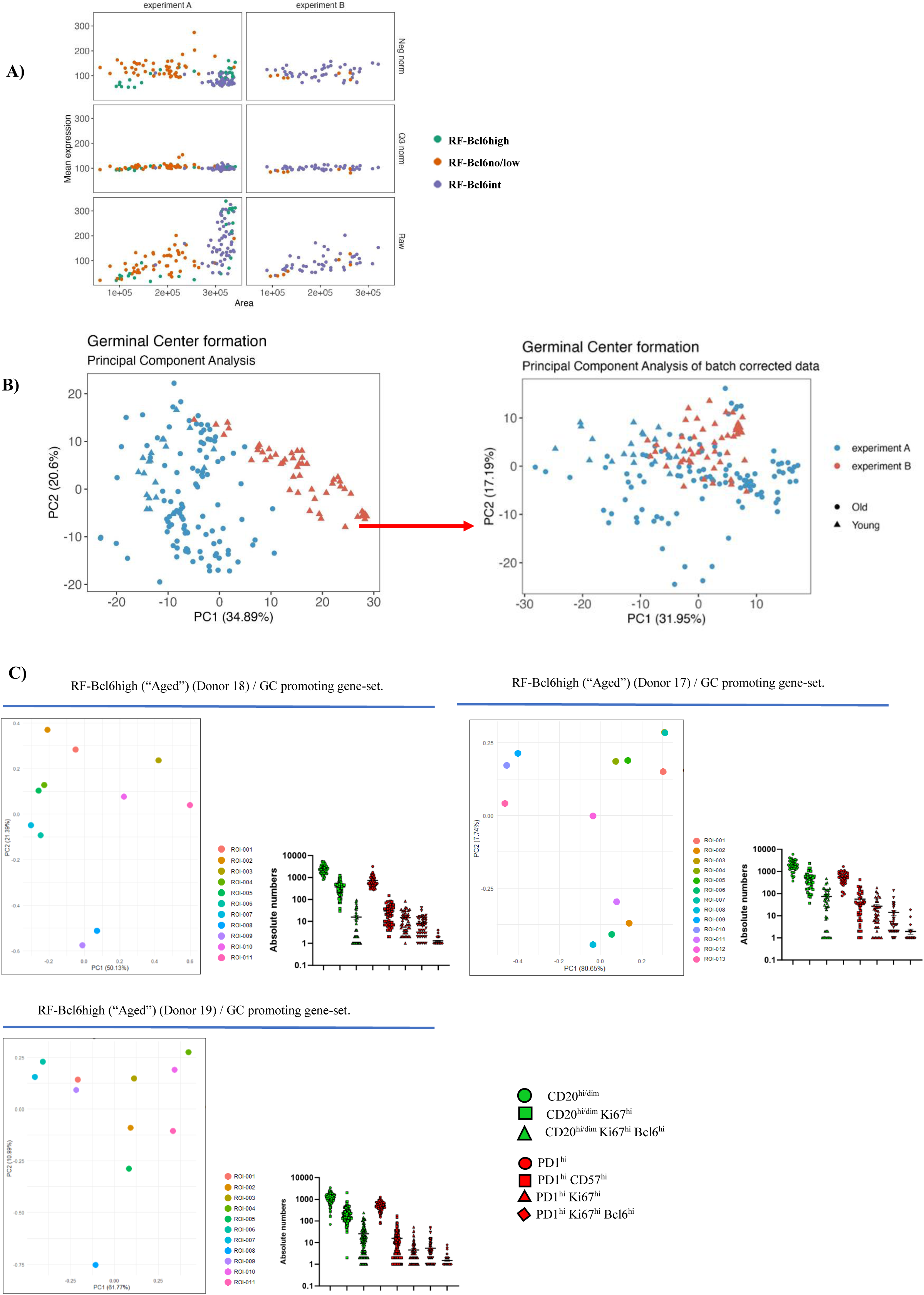
**A)** Visualizing plots of the normalized GeoMx-generated data per ROI. The figure represents a series of scatter plots comparing the mean expression levels across different ROIs for two experiments (A and B) under three different normalization conditions: Negative normalization (Neg norm), Q3 normalization (Q3 norm), and Raw data (Raw). The data points are color-coded to represent the three different groups: RF-Bcl6high (green), RF-Bcl6int (orange) and RF-Bcl6no/low (purple). **B)** Principal Component Analysis (PCA) plots before and after batch correction. The first plot illustrates the PCA before batch correction (left image), with data from Experiment A (cyan) and Experiment B (red) showing clear separation along PC1 (34.89%) and PC2 (20.6%), indicating batch effect. The second plot (right image), after batch correction using the Trimmed Mean of M-values (TMM) normalization and RUV-4 correction, displays data from Experiment A (cyan) and Experiment B (red) along PC1 (31.95%) and PC2 (17.19%). **C)** PCA plots showing the distribution of the follicular areas (ROIs), based on the expression of the gene set favoring GC-development, in RF-Bcl6high subgroup (Donor 18, Donor 17, Donor 19). The plots were generated using RStudio based on genes expression extracted from the GeoMx. The absolute numbers of B (CD20^high/dim^, CD20^high/dim^Ki67^high^, and CD20^high/dim^Ki67^high^Bcl6^high^) and TFH (PD1^high^, PD1^high^CD57^high^, PD1^high^Ki67^high^, PD1^high^Ki67^high^Bcl6^high^) subsets in individual follicles from the same individuals are shown too.

**Suppl. Figure 4.**
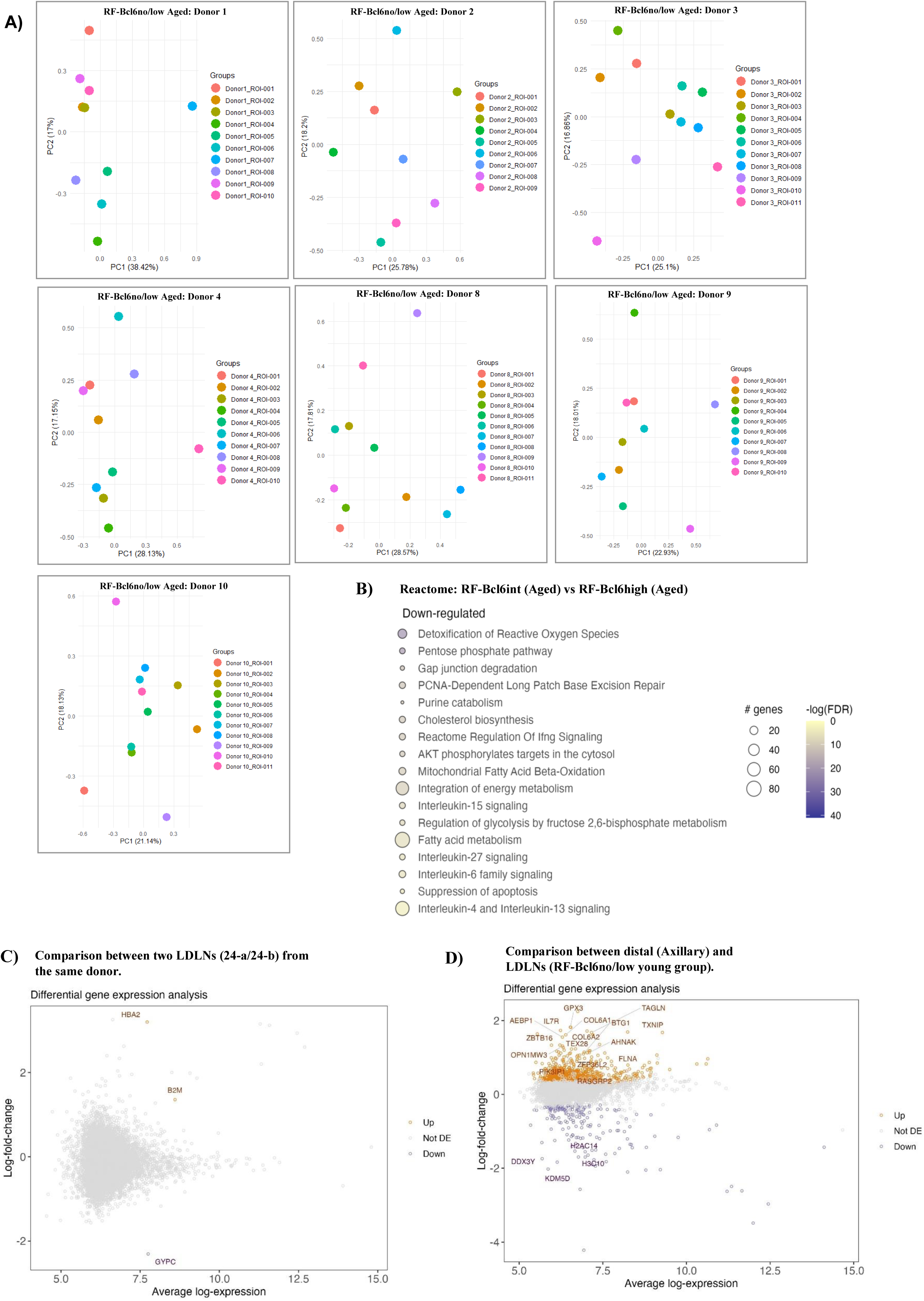
**A)** PCA plots showing the distribution of follicular areas (ROIs), based on the expression of the gene set favoring TH1 responses, in the RF-Bcl6no/low group (Donor 1, Donor 2, Donor 3, Donor 4, Donor 8, Donor 9, Donor 10). The plots were generated using RStudio based on genes expression extracted from the GeoMx. **B)** Reactome pathways analysis showing pathways upregulated in “Aged” RF-Bcl6high compared to “Aged” RF-Bcl6int (downregulated). **C)** Volcano plot of gene expression for two RF-Bcl6no/low LD-LNs (24-a and 24-b) from the same donor (Donor 24). **D)** Volcano plot of gene expression between distal (Axillary, Donor 25) RF-Bcl6no/low and LD-LNs RF-Bcl6no/low from the “Young” group. Volcano plot is depicted with the log-fold change of each gene and the average log expression of each gene. Orange dots show genes significantly increased in the Axillary LN and purple dots genes significantly increased in the LD-LNs from the same RF-Bcl6no/low “Young” group.

## Acknowledgements

The authors would like to thank Dr Natalie Piazzon (operational director of the Tissue Biobank), Damien Maison and Emilie Lingre, Institute of Pathology, CHUV, for their help with the tissue processing.

## Authorship Contributions

C.B and K.I performed experiments and image analysis, S.G, and M.O contributed to image analysis, C.B and S.G drafted the manuscripts, M.B and R.G. performed and supervised the spatial transcriptomic analysis, C.B. participated in the transcriptomic analysis, S.B. and C.B. performed neighboring analysis, O.Y.C. supervised the statistical analysis, J.B supervised the SQL clustering analysis, M.S. and S.B. assisted with the clinical data, serological measurements and pathological evaluation/information, N.S. provided serological measurements, M.J.F, J.W.L. and L.de.L, provided tissue material and pathological information, G.P. assisted with the interpretation of the immunological data and the manuscript preparation, CP conceived, designed and supervised the study, interpreted data, and edited the manuscript. All authors have read, edited and approved the final version for submission.

## Data sharing statement

The authors agree to share all publication-related data. For further information, please contact the corresponding author at.

